# Netrin4 is a new target specific factor, ensuring adult sympathetic neuron survival via promoting protein synthesis

**DOI:** 10.1101/2023.12.23.573195

**Authors:** Zhu Zhu, Lili Zhou, Yuanjiao Wang, Xiaoxuan Zhang, Dongdong Zhang, Xuzhao Li, Bingrui Zhao, Jie-Min Jia

## Abstract

How mature neurons survive under homeostasis is a question of utmost importance and is known to be different from developing neurons. However, the understanding of this regard remains largely unknown. Here, based on the relationship between the sympathetic cervical ganglia (SCG) and the arterial networks of projecting and targeting organs, we report that the secretome of cerebral, but not peripheral, arterial smooth muscles (SMC) was required for the survival of sympathetic neurons. Among the secretome, we further identified that netrin-4, encoded by the ntn-4 gene, only entered neurons and not glia and played a crucial role both in vitro and in vivo. This was demonstrated with three independent lines of tamoxifen-inducible SMC-specific conditional knockout mice (cKO). Notably, in cKO mice, the local supply of exogenous netrin-4 confined to SCG selectively rescued neuronal necroptosis, which otherwise consistently occurred in a specific subgroup of SCG neurons that innervate cerebral SMCs. Mechanistically, we demonstrated that cerebral netrin-4 was endocytosed at the neurovascular interface and retrogradely long transported to peripheral soma in SCG, where it differentially regulated mRNA translations. This regulation suppressed vacuolization and neuronal necrosis, both of which took place spontaneously in cKO mice. The former is immediately followed by the latter when we injured axons using two-photon laser ablation. The findings revealed a new principle of neurovascular interactions vital for mature neuron survival, implying that under circumstances of cerebral SMC insufficient secretion, such as natural aging, may initiate mature neuronal loss due to uncontrolled vacuolization.

---

The lifelong survival of mature neurons is a fundamental requirement for ensuring various bodily functions throughout an individual’s life[1]. During embryonic development, only half of the neurons formed undergo a competitive process, relying on limited neurotrophic factors secreted by target organs to initiate anti-apoptotic programs, allowing them to mature and survive[2,3]. In contrast to the distinct process observed during embryonic development, mature neurons in adulthood do not readily enter apoptotic pathways, maintaining a stable population throughout an individual’s lifespan[4].

However, the long-term survival of mature neurons faces challenges during periods of neurodegeneration, where various neurodegenerative diseases lead to functional impairment or even loss of adult neurons[5–7]. Therefore, understanding the survival mechanisms of mature neurons is not only crucial for comprehending how neurons persist over the long term but also offers insights into how various neurodegenerative diseases ultimately result in neuronal death. However, the mechanisms underlying the survival of mature neurons remain unclear to date, necessitating substantial research efforts to unravel this mystery.

The system focusing on the Superior Cervical Ganglion (SCG) has long been a well-known paradigm in the field for studying the mechanisms of neuronal death and survival. This system has provided valuable insights into understanding the survival mechanisms of central neurons[8]. The target organ of the SCG is the cerebral vasculature, and abnormalities in the sympathetic nervous system are closely linked to cerebrovascular diseases[9,10]. However, the survival mechanisms of mature SCG neurons and their relationship with cerebral vasculature remain elusive.

The output fibers of mature SCG neurons accompany the internal carotid artery into the cranium, innervating intracranial meningeal arteries[11,12]. Research has reported that arterial smooth muscle cells (aSMCs) derived from the aorta promote the in vitro survival of newborn SCG neurons[13,14]. However, whether aSMCs secrete neurotrophic factors to sustain the long-term survival of mature neurons has not been investigated.

In addition, previous studies have shown that mature SCG neurons in in vitro cultures can resist deprivation of nerve growth factor (NGF) and survive[15,16]. However, at the systemic level, whether mature SCG neurons depend on the secretion of another novel neurotrophic factor from target cells for their survival remains to be explored.

Our preliminary exploratory work indicated that, through an in vitro cell culture research system, the “secreted protein mixture” from human-derived blood vessel smooth muscle cells (HBvSMCs) and primary mouse meningeal aSMCs has a nutritive effect on mouse hippocampal and cortical neurons. This is manifested by effectively promoting the developmental processes of neuronal dendrites and increasing neuronal survival rates [17].

In this study, we further discovered that the secreted protein mixture from aSMCs not only enhances the survival rate of primary cells derived from immature SCG neurons of neonatal mice but also increases the survival rate of primary cells from mature SCG neurons of adult mice by inhibiting the cell necroptosis. Following screening and validation, Netrin4 emerged as a potential key neuroprotective factor secreted by aSMCs to sustain the survival of adult neurons.

We confirmed that conditional knockout of Netrin4 in adult aSMCs led to neuronal death, vacuolization, and loss in the SCG. Local supplementation of Netrin4 in the SCG was able to rescue these pathological changes. We also revealed that Netrin4, upon internalization from the distal ends of neurons and retrograde transport to the cell body, may play a crucial role in sustaining specific protein synthesis, representing a key mechanism for maintaining the survival of mature neurons.

This article employs cell-specific gene intervention techniques to elucidate the nutritional relationship between the long-term survival of adult SCG neurons and the secreted proteins from cerebral aSMCs. The aim is to decipher the fundamental principles allowing neurons to age synchronously with the organism. The findings seek to provide a theoretical basis for the future development of preventive strategies against neuronal death during the aging process, specifically triggered by insufficient secretion capacity in arterial cells.

## Methods

### Animals

Mice, including C57BL/6J mice and transgenic lines such as THCre:Ai14, THCreER:Ai47, SMACreER:Ai14, PDGFRβCreER, Myh11CreER, and Rosa26-floxp-Netrin4 mice were employed and housed at Animal Experimental Center of West Lake University. The study received ethical approval, and mice were raised in a specific pathogen-free (SPF) environment to ensure experimental integrity. Stringent adherence to animal welfare guidelines was maintained throughout the experimental procedures to guarantee ethical and humane treatment of the mice.

### cardiac perfusion and cryosection

Cardiac perfusion and cryosectioning were conducted for subsequent molecular and histological analyses. For cardiac perfusion, mice were humanely sacrificed after being anesthetized with a 1% pentobarbital sodium solution. Their hearts were rapidly excised and immersed in PBS to ensure thorough blood clearance. A syringe pump was employed for aortic root perfusion with PBS until complete vascular clearance was achieved. Subsequently, 4% paraformaldehyde was perfused.

In the cryosectioning process, the superior cervical ganglion (SCG) tissues were retrieved and immersed in a 30% sucrose solution for 6 hours to ensure optimal cryoprotection. Subsequently, the tissues were frozen and sectioned to a thickness of 8 micrometers using a cryostat (3050S, Leica, Germany).

### Medullary cisterna injection

Mice were anesthetized with 2% isoflurane, and a stereotactic setup was utilized for stability. A small craniotomy exposed the cisterna magna, into which a 30G niddle was gently inserted for 10 ul tracer injection. The infusion speed 2 ul/ min, regulated by a microsyringe pump, occurred with careful monitoring to avoid complications.

### Immunofluorescence

Harvested tissues, including vessels and frozen sections, were washed with PBS, followed by blocking and permeabilization steps. Primary antibody incubation occurred overnight at 4℃, followed by washing and incubation with a fluorescently labeled secondary antibody. Nuclei were optionally stained with DAPI, and tissues were mounted for imaging using a fluorescence microscope (ZSM900, Zeiss, Germany). Quantification was performed using image analysis software Zen for data interpretation.

### SCG primary cell culture

SCG were dissected, and neurons were enzymatically digested using 1 mg/ml Collagenase D at 37°C for 30 minutes. The obtained cell suspension was cultured in Neurobasal-A medium supplemented with B27 and GlutaMAX for optimal growth conditions. Cultures were maintained in a humidified incubator at 37℃ with 5% CO_2_.

### Western Blotting

Cell or tissue samples were homogenized and proteins extracted using RIPA (Thermofisher, USA). After quantification, equal protein amounts were loaded onto a gel for SDS-PAGE. Proteins were then transferred to a PVDF membrane, blocked to prevent nonspecific binding, and probed with primary and secondary antibodies. Enhanced chemiluminescence was applied for band visualization, and signals were captured using a chemiluminescence detection system.

### RT-PCR

Gene expression in tissues and cells was assessed using RT-PCR. Total RNA extraction and cDNA synthesis was performed with the High-Capacity cDNA Reverse Transcription Kit. Specific primers for 18S, Netrin4, ACTA2, and CD31 were used. Data, normalized to the internal reference gene 18S, provided insights into gene expression profiles in the studied samples.

### Laser Speckle Contrast Imaging (LSCI)

LSCI was employed for real-time observation of cerebral blood flow (CBF) patterns in awake mice. Prior to experimentation, animals were acclimated to the experimental setup for one hour daily over three consecutive days using a mouse fixation device. Subsequently, under continuous isoflurane anesthesia, the animals underwent head skin incision for laser speckle observation. Post-surgery, daily subcutaneous injections of meloxicam were administered. Experimental conditions were optimized to minimize external light interference, and stable room temperature was maintained.

### Detection of Protein Synthesis

To evaluate protein synthesis, O-propargyl-puromycin (OPP) was utilized in both cell culture and animal experiments. In cell culture, 20 uM OPP was added to the supernatant (1:1000 dilution), followed by a 30-minute incubation at 37℃. After fixing cells with 4% PFA, OPP incorporation was detected using a click chemistry assay. In vivo, each mouse received a 30 uL injection of 20 uM OPP. Following one hour of homecage housing, animals underwent heart perfusion, and OPP signal detection was carried out on frozen tissue sections through a click reaction. For combined click reaction and immunofluorescence staining, it is generally conducted to perform the click reaction first.

### Statistical Analysis and Data Visualization

Statistical analyses were employed to assess the significance of data differences among groups. T-tests were employed for two-group comparisons, assuming normal distribution post normality test verification. ANOVA was utilized for comparisons involving three or more groups, allowing the identification of significant differences. Immunofluorescence intensity and colocalization analyses were performed using Zen and Imaris 9.3 software. Graphical representation of the data was achieved using GraphPad and the R language ggplot2 package.

## Results

### 1 postganglionic fiber of SCG innervate the intracranial ICA

In order to delineate the pathway of postganglionic fibers from the SCG into the cranium, we perfused the hearts of mice with pink latex, followed by dissection along the CCA to isolate the ICA up to the carotid foramen, exposing the skull base and its major branches (**Fig 1**, **A**). To distinguish between intracranial and extracranial structures, we further injected Dextran-FITC into the cerebellomedullary cistern of THCreER:Ai14 mice and found that the intracranial blood vessels were filled with FITC fluorescence (**Fig S1, B**). Postganglionic nerve fibers from the SCG formed bundled structures wrapping around the extracranial ICA (exICA) and extending into the cranium, displaying a plexiform arrangement after passing through the carotid foramen, surrounding the intracranial ICA (inICA) and its branches (**Fig 1**, **B)**. Imaging after the sequential dissection from CCA to inICA validated these structural features (**Fig S1, A**). By immunofluorescent staining of total nerve endings (Tuj1) and sympathetic nerve endings (TH) after dissecting inICA and exICA, we found that approximately 17% of the nerve endings innervating exICA were sympathetic, while approximately 84% of the nerve endings innervating inICA were sympathetic (**Fig 1**, **C-E**). To accurately define the structural zones of the SCG based on the distribution of blood vessels, we observed SMACreER:Ai14 mice at the bifurcation of CCA and SCG. We discovered that the primary blood supply arteries to the SCG originated from branches of the external carotid artery (ECA), specifically the lingual artery. Immediately upon branching, the lingual artery split into two branches, both entering the SCG at the external carotid nerve (ECN) portion. The upper branch formed three main branches, supplying the upper and middle portions of the SCG, while the lower branch was responsible for supplying the ST segment. The upper branch immediately split into three smaller branches upon entering ECN, with the first two supplying the upper portion of the SCG and the third supplying the middle portion. Drawing an extension line from the base of the vessel at the root of the second branch, the portion above is defined as the internal carotid nerve (ICN) portion; and drawing an extension line from the lower edge of ECN serves as the boundary between ECN and ST portions (**Fig 1**, **F**; **Fig S1, C**). To clarify the distribution characteristics of neuronal cell bodies responsible for intracranial innervation within the SCG, we injected the retrograde tracer WGA into the CSF. Continuous observations at 12 hours, 24 hours, and 48 hours after injection revealed a continuous increase in the number of labeled neurons within the SCG. After 48 hours of retrograde tracing, approximately 15% of SCG neurons were labeled, with 92% of labeled neurons located in the ICN portion (**Fig 1** **G-I**).

**Figure 1.**
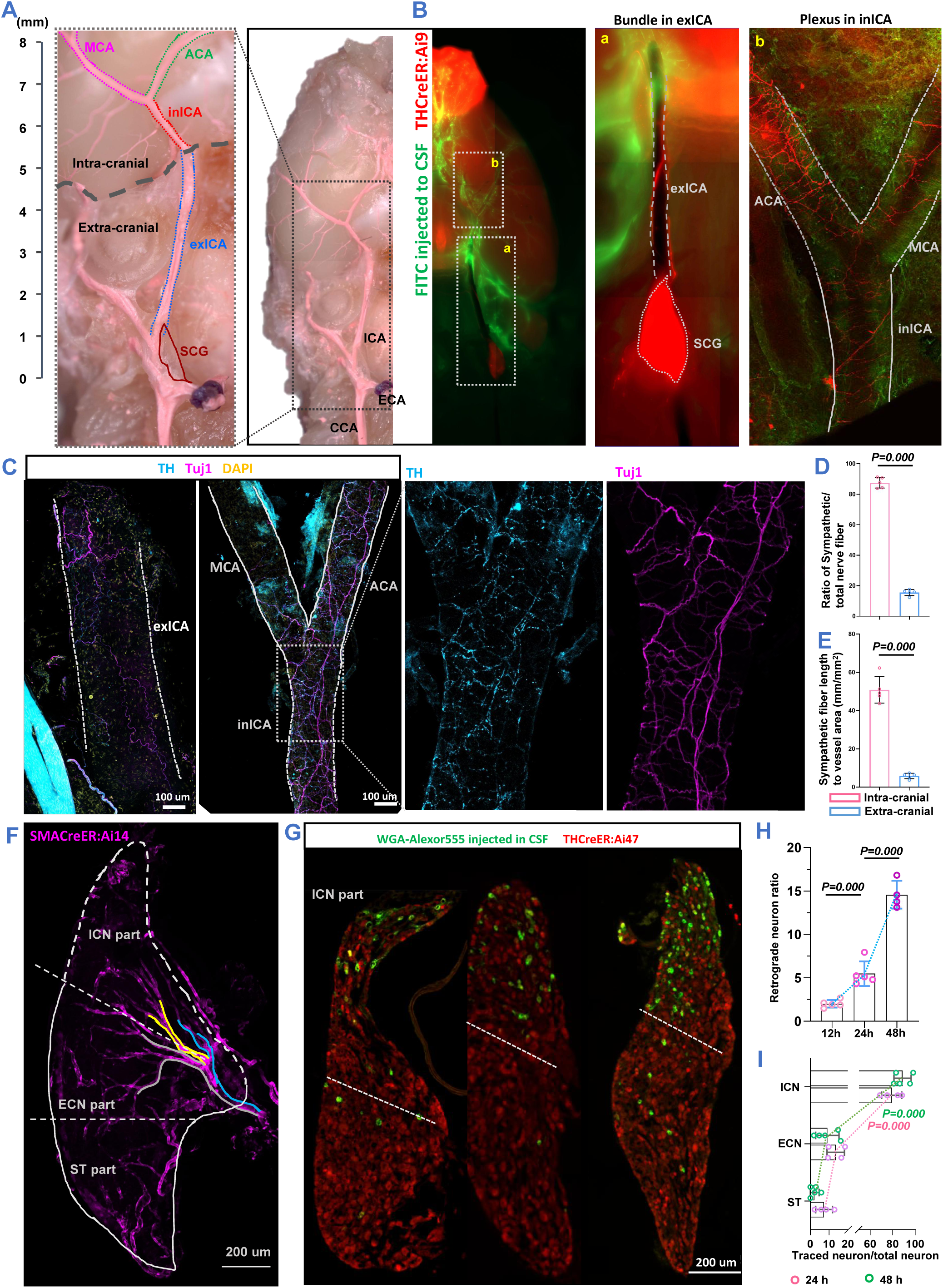
postganglionic fiber of SCG innervate the intracranial ICA and its major branches. **A)** The right panel depicts the ventral aspect of the mouse skull base anatomy, where the CCA branches into the ECA and ICA, with the SCG located deep at the vascular bifurcation. The dashed line in the left panel indicates the intracranial extension of the ICA, traversing through the carotid foramen at the skull base, continuing as the inICA on the brain surface, and further branching into the MCA and ACA. The gray dashed lines delineate the boundaries between intracranial and extracranial regions. **B)** Dextran-FITC was injected into the CSF to label the intracranial region, and imaging of the SCG and postganglionic nerve fiber distribution was conducted using THCreER: Ai14 mice. The experiment revealed that postganglionic fibers from the SCG extended in bundles along the exICA, entered the skull at the carotid foramen, and formed a plexus surrounding the inICA and its branches. **C, D, E)** Stripping of exICA and inICA was performed for immunofluorescent staining with TH, Tuj1, and DAPI, marking sympathetic nerve terminals, neuronal terminals, and cell nuclei, respectively. Statistical analysis revealed that approximately 84% of nerve terminals on the surface of inICA were sympathetic terminals, while sympathetic nerve terminals accounted for approximately 17% on the surface of exICA. Moreover, the length of sympathetic nerve fibers distributed on the surface of inICA was significantly higher than that on exICA. **F)** After tissue clearing of SCG from SMACreER:Ai14 mice, imaging was conducted to observe the vascular distribution in the SCG. The arteries supplying blood to the SCG entered from the ECN end, stably presenting three branches, delineated in blue, yellow, and gray to represent the upper, middle, and lower branches, respectively. A prolongation line was drawn along the base of the middle branch to define the ICN portion. Another vertical line was drawn through the lower end of ECN perpendicular to the longitudinal axis of the SCG, distinguishing the ECN portion from the ST portion. **G, H, I)** After intracisternal injection of WGA into the medullary cistern, the distribution of retrogradely traced SCG neurons was observed at 12, 24, and 48 hours. The region above the dashed line represents the ICA portion. Approximately 15% of SCG neurons were retrogradely labeled 48 hours after WGA CSF injection, and around 90% of these neurons were located in the ICN portion. CCA: common carotid artery; ECA: external carotid artery; ICA: internal carotid artery; SCG: superior cervical ganglion; inICA: intracranial ICA; exICA: extracranial ICA; MCA: middle cerebral artery; ACA: anterior cerebral artery; CSF: cerebrospinal fluid; ICN: internal carotid nerve; ECN: external carotid nerve; ST: sympathetic trunk.

### 2 Secreted supernatant of intracranial vSMCs maintained adult neurons survival

In order to investigate the impact of intracranial vSMCs on neuronal survival, adult SCG neurons were isolated and cultured (**Fig S2, A, B**). Compared to the Neurobasal-A culture group and the NGF treatment group, seceted supernatant from HBvSMCs or primary isolated mouse brain vSMCs effectively increased the density of neurons in vitro from 12 to 96 hours (*P<0.05*, **Fig 2**, **B; Fig S2, F**). The 48-hour survival density of neurons treated with 0.5X to 2X vSMCs seceted supernatant showed a dose-response relationship (R=0.76, *P<0.05*). However, seceted supernatant from heat-inactivated vSMCs had no effect on neuronal survival (**Fig 2**, **A**). When cell death inhibitors were added to the basic neuron culture medium, it was found that, compared to the Neurobasal-A culture group, necroptosis inhibitor Nec-1, pyroptosis inhibitor VRT-043198, and ferroptosis inhibitor Fer-1 added effectively increased the survival density of adult neurons (*P<0.05*), while apoptosis inhibitor Z-VAD-fmk was ineffective. Adding cell death inhibitors to vSMCs seceted supernatant revealed that, compared to the vSMCs seceted supernatant culture group, VRT-043198 and Fer-1 added effectively increased the survival density of adult neurons (*P<0.05*), while necroptosis inhibitor Nec-1 was ineffective. This suggests that vSMCs may enhance neuronal survival by inhibiting cell necroptosis (**Fig 2**, **E-G**).

**Figure 2.**
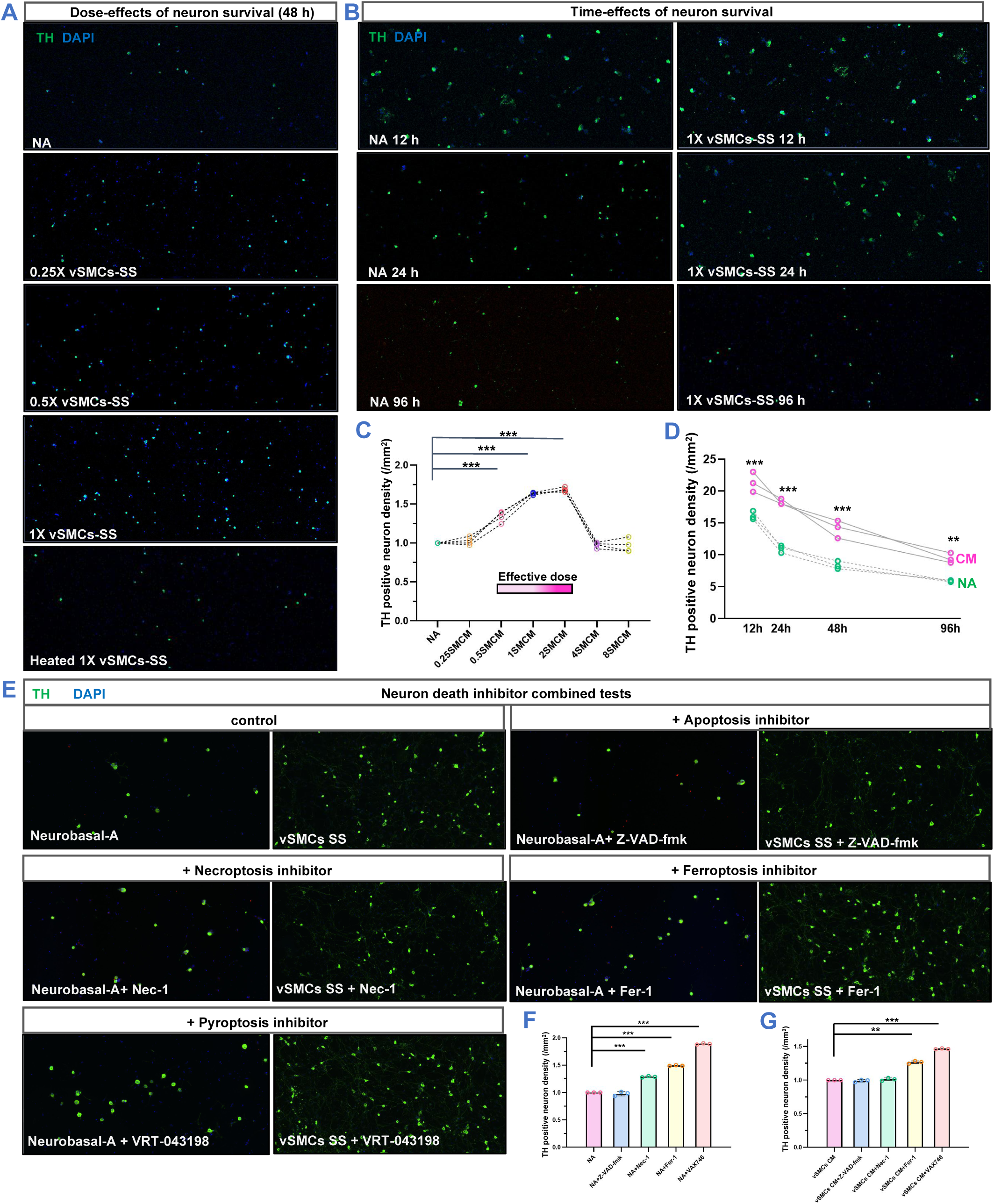
The secreted supernatant derived from intracranial vascular smooth muscle cells (vSMCs) enhance the survival of adult SCG neurons by suppressing cell necroptosis. **A, C)** When culturing adult SCG neurons with secreted supernatant from vSMCs at different concentrations, it was observed that concentrations ranging from 0.5X to 2X significantly enhanced the survival of adult SCG neurons. The secreted supernatant from vSMCs subjected to heat inactivation did not lead to an increase in the survival rate of adult SCG neurons. **B, D)** Utilizing the conditioned medium from vSMCs for the in vitro culture of adult SCG neurons effectively promoted neuron survival within the first 12 to 96 hours of isolation and culture. **E, F, G)** Experimental findings from co-treating adult SCG neurons with cell death inhibitors revealed that the use of apoptosis inhibitors did not elevate the survival rate of adult SCG neurons. Additionally, necroptosis inhibitors did not further improve the survival rate of neurons cultured with vSMCs-secreted supernatant.

### 3 The brain vascular-specific Netrin4 sustain the survival of sympathetic neurons

Published single-cell sequencing data revealed that, compared to cerebral vascular endothelial cells and pericytes, cerebral vSMCs highly express 42 genes coding secreted proteins (**Fig 3 A**). Through the isolation of adult SMACreER:Ai14 mouse brain vSMCs and other cell types, followed by transcriptome analysis, it was revealed that vSMCs exhibit higher expression of 30 genes encoding secreted proteins compared to glial cells (**Fig 3 B**). Following the individual neutralization of Netrin4 within the secreted supernatant of cerebral vSMCs using antibodies, it was noted that the survival density of adult SCG neurons cultured in the supernatant supplemented with Netrin4 antibody exhibited a significant decrease compared to the vSMCs supernatant culture group (*P<0.05*), yet remained higher than that in the Neurobasal-A culture group (*P<0.05*). However, neutralizing GDF-15 in the vSMCs secreted supernatant with antibodies did not impact the neuronal survival density (**Fig S3**). The inclusion of Netrin4 in the basal culture medium for neurons significantly augmented both the survival density and size of adult superior cervical ganglion (SCG) neurons. The effective concentration was determined to be 25 ng/ml, with the optimal concentration identified as 100 ng/ml (**Fig 3** **C-E**). Upon incorporating Netrin4 into the basal culture medium of adult SCG neurons, immunoblot experiments revealed that Netrin4 significantly amplifies the expression levels of neuronal ERK, AKT, and pS6 (**Fig 3** **F, G**). This observation implies that Netrin4 increases the protein synthesis levels in adult SCG neurons.

**Figure 3.**
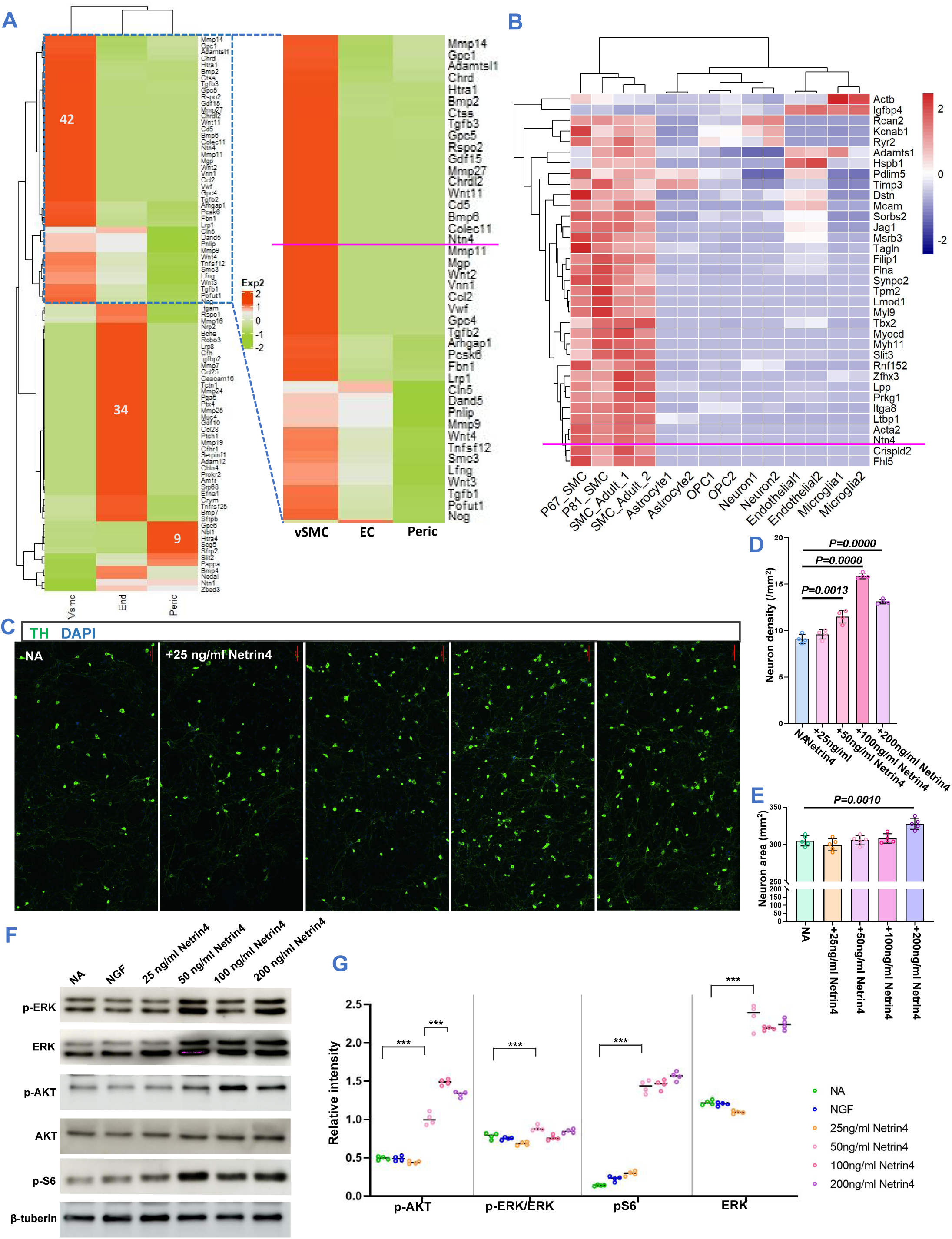
Netrin4 may be one of the neurotrophic factors secreted by cerebral vSMCs. **A)** Reanalysis of previous single-cell sequencing data revealed that, compared to cerebral vascular endothelial cells and pericytes, cerebral vSMCs possess secretory activity, capable of secreting 42 relatively specific secreted proteins. **B)** Transcriptome analysis, through the sorting of cerebral vSMCs and common glial cells in the brain, revealed that adult cerebral vSMCs can secrete specific proteins. **C-E)** The addition of Netrin4 to the neuronal basal culture medium effectively increased the survival density and neuron size of adult SCG neurons, with the effective concentration being 25 ng/ml and the optimal concentration being 100 ng/ml. **F, G)** Upon the addition of Netrin4 to the basal culture medium of adult SCG neurons, immunoblot experiments demonstrated that Netrin4 effectively enhances the expression levels of neuronal ERK, AKT, and pS6, suggesting that Netrin4 elevates the protein synthesis levels in adult SCG neurons. NA: Neurobasal-A.

The intracranial segments of the CCA and ICA, as well as the intracranial branches of the ICA on the brain ventral side, were dissected. Immunofluorescence staining revealed higher expression of Netrin4 in the inICA compared to the exICA (*P<0.05,* **Fig 4** **A, B**). Cross-sectional slices of intracranial and extracranial arteries underwent immunofluorescence staining, elucidating that Netrin4 exhibits a predominant distribution in intracranial arteries. Its primary expression is observed in vSMCs and the basement membrane, with no discernible Netrin4 signal detected in endothelial cells, as indicated by CD31 staining (**Fig S4, A-C**).Using SMACreER:Ai14 mice, vSMCs were sorted from different arteries or tissues inside and outside the skull, and RT-PCR analysis showed that the vSMCs of the inICA and MCA in the meningeal arteries exhibited high expression of Netrin4 mRNA. Netrin4 mRNA expression was almost undetectable in vSMCs derived from the exICA and SCG. Arterial vSMCs showed higher expression of Netrin4 mRNA compared to other cells from vessels or tissues (*P<0.05,* **Fig 4** **C-F**). Immunoblot analysis of Netrin4 expression in different intracranial and extracranial vessels or tissues revealed high expression of Netrin4 in the intracranial segments, namely inICA and MCA (**Fig 4** **G, H**).

**Figure 4.**
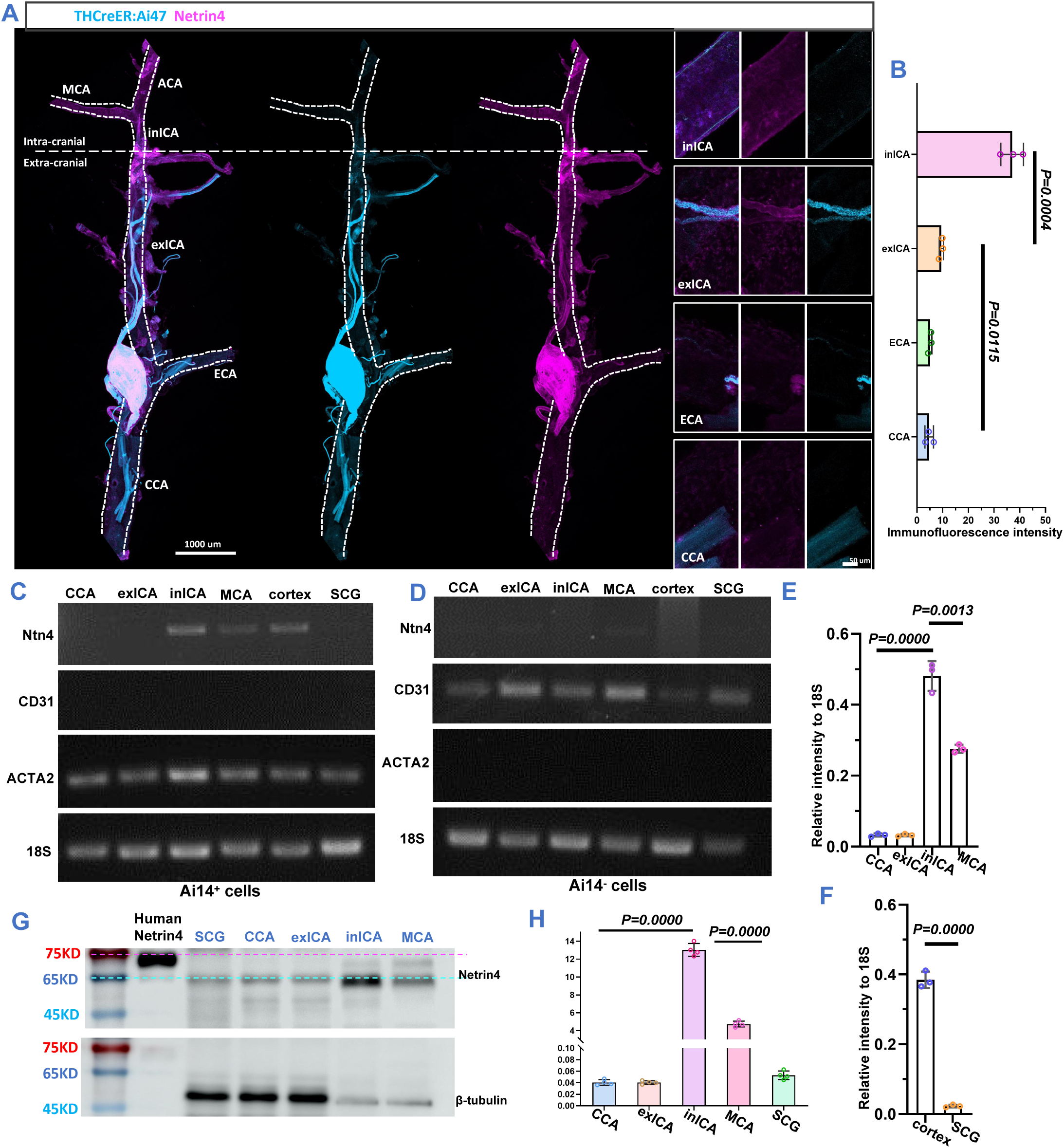
Netrin4 is highly expressed in intracranial arterial SMCs and can promote the survival of adult SCG neurons. **A, B)** The intracranial segments of the CCA and ICA, as well as the intracranial branches of the ICA on the brain ventral side, were dissected. Immunofluorescence staining revealed higher expression of Netrin4 in the inICA compared to the exICA. **C, D, E, F)** VSMCs were sorted from different arteries or tissues inside and outside the skull, and RT-PCR analysis showed that the vSMCs of the inICA and MCA in the meningeal arteries exhibited high expression of Netrin4 mRNA. Netrin4 mRNA expression was almost undetectable in vSMCs derived from the exICA and SCG. Arterial vSMCs showed higher expression of Netrin4 mRNA compared to other cells from vessels or tissues. **G, H)** Immunoblot analysis of Netrin4 expression in different intracranial and extracranial vessels or tissues revealed high expression of Netrin4 in the intracranial segments, namely inICA and MCA.

From birth to 130 days, the size of the SCG gradually increases, reaching and maintaining its maximum from 130 to 180 days. However, from 230 to 270 days after birth, the SCG size significantly decreases compared to the 130-day mark (**Fig 5** **A**; **Fig S5 A, B**). Utilizing SMACreER:Netrin4^-/-^ mice with a conditional knockout of arterial vSMC Netrin4 and analyzing by landmarker identity, notable changes in SCG morphology were observed compared to heterozygous and wild-type mice, primarily characterized by a reduction in the ICN segment (**Fig 5** **B, C; Fig S5 C**). On the 7th day of conditional knockout of arterial vSMC Netrin4 in SMACreER:Netrin4^-/-^ mice aged 130 days post-birth, the SCG volume was significantly smaller than in wild-type mice (*P<0.05*). Additionally, using PDGFRβCreER:Netrin4^-/-^ and Myh11CreER:Netrin4^-/-^ mice with conditional knockout of vascular mural cell Netrin4 on the 7th day, a significant reduction in SCG volume was observed compared to wild-type mice (*P<0.05,* **Fig 5**, **D**). Further analysis of SCG morphology revealed that on the 5th day of conditional knockout of arterial SMC Netrin4 in SMACreER:Netrin4^-/-^ mice, the proportion of ICN terminal volume to the total SCG volume significantly decreased, suggesting an initial reduction in ICN terminal volume (**Fig 5**, **E**). Slicing the SCG of Netrin4 conditional knockout mice and counting the total number of neurons revealed a significant decrease in the total number of neurons and glial cells in both heterozygous and homozygous SMACreER:Netrin4 conditional knockout (CKO) mice (*P<0.05*), with a more pronounced decrease in the total number of neurons (**Fig 5**, **F, I, J**). Heterozygous and homozygous SMACreER:Netrin4 CKO mice both showed a decrease in cerebrospinal fluid NE levels, while an increase in SCG NE levels was observed only in homozygous SMACreER:Netrin4 CKO mice (**Fig 5**, **G, H**). Immunofluorescence staining in homozygous SMACreER:Netrin4 CKO mice detected the necroptosis marker pMLKL expression in both SCG neurons and glial cells, while the apoptosis marker cleaved-Caspase3 was not detected (**Fig 5**, **K-O**).

**Figure 5.**
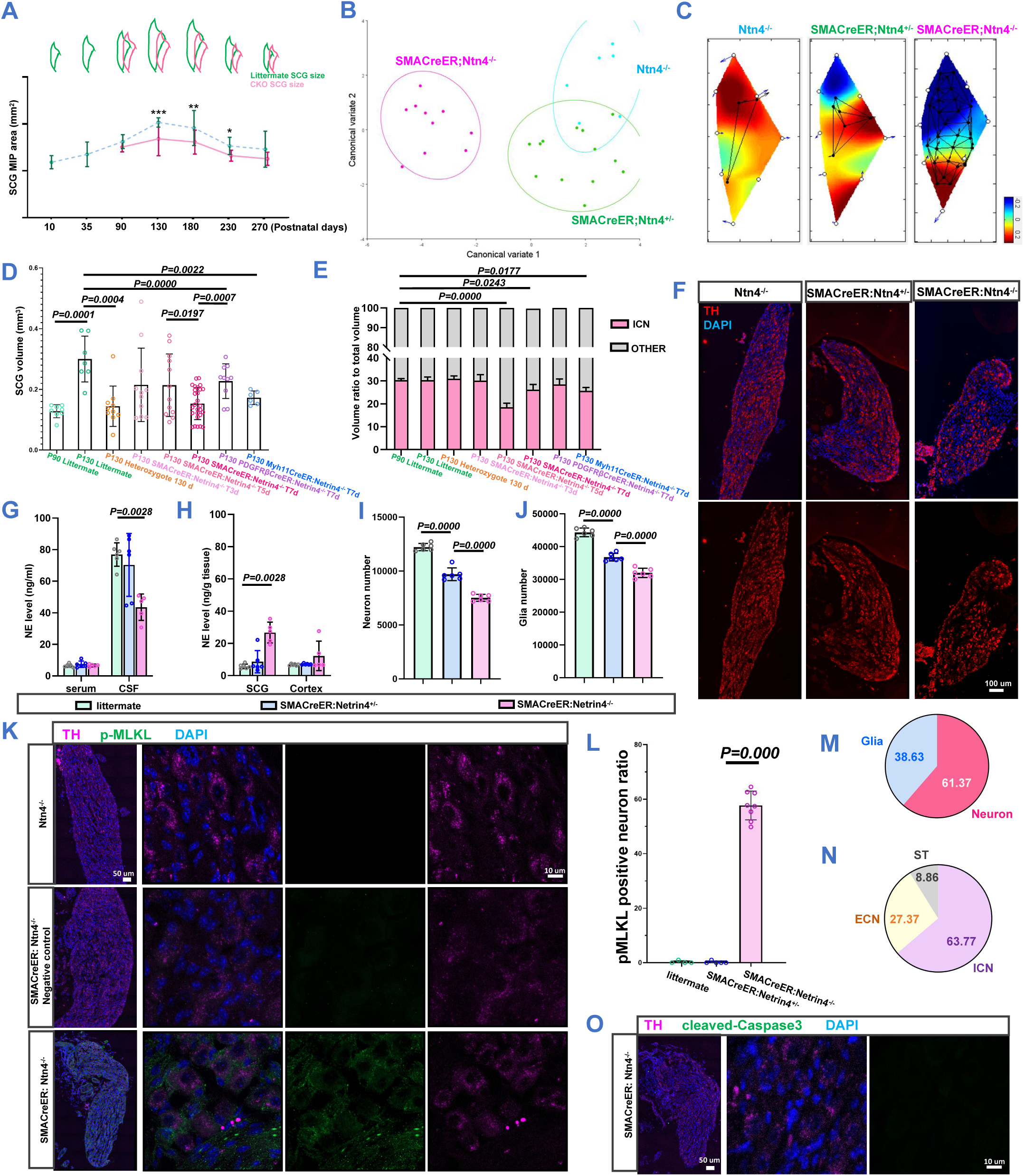
Conditional knockout of vSMCs-derived Netrin4 leads to necroptosis and loss of adult mouse SCG neurons. **A)** From birth to 130 days, the size of the SCG gradually increases, reaching and maintaining its maximum from 130 to 180 days. However, from 230 to 270 days after birth, the SCG size significantly decreases compared to the 130-day mark. **B, C)** Using SMACreER:Netrin4^-/-^ mice with conditional knockout of arterial SMCs Netrin4, it was observed that the morphology of SCG changed significantly compared to heterozygous and wild-type mice, primarily characterized by a reduction in the ICN segment. **D)** On the 7th day of conditional knockout of arterial SMCs Netrin4 in SMACreER:Netrin4^-/-^ mice aged 130 days post-birth, the SCG volume was significantly smaller than in wild-type mice. Additionally, using PDGFRβCreER:Netrin4^-/-^ and Myh11CreER:Netrin4^-/-^ mice with conditional knockout of vascular mural cell Netrin4 on the 7th day, a significant reduction in SCG volume was observed compared to wild-type mice. **E)** Further analysis of SCG morphology revealed that on the 5th day of conditional knockout of arterial SMC Netrin4 in SMACreER:Netrin4^-/-^ mice, the proportion of ICN terminal volume to the total SCG volume significantly decreased, suggesting an initial reduction in ICN terminal volume. **F, I, J)** Slicing the SCG of Netrin4 conditional knockout mice and counting the total number of neurons revealed a significant decrease in the total number of neurons and glial cells in both heterozygous and homozygous SMACreER:Netrin4 CKO mice, with a more pronounced decrease in the total number of neurons. **G, H)** Heterozygous and homozygous SMACreER:Netrin4 CKO mice both showed a decrease in cerebrospinal fluid NE levels, while an increase in SCG NE levels was observed only in homozygous SMACreER:Netrin4 CKO mice. **K, L, M, N, O)** Immunofluorescence staining in homozygous SMACreER:Netrin4 CKO mice detected the necroptosis marker pMLKL expression in both SCG neurons and glial cells, while the apoptosis marker cleaved-Caspase3 was not detected.

Utilizing retrograde labeling through the medullary cistern in SMACreER:Ai14 mice, WGA-Alexor647 was introduced into the CSF to trace neurons innervating the intracranial structures, while WGA-Alexor488 was injected into the salivary gland to trace neurons innervating peripheral regions. Neuronal fluorescence signals were reconstructed as spots and quantified (**Fig 6** **A-C**). The experiment unveiled that, in comparison to littermate control mice, the number of neurons in the ICN portion in SMACreER:Netrin4^-/-^ mice at 130 days post-birth significantly decreased at CKO 5 days and 7 days (*P<0.05*). This observation was also noted in heterozygotes and two other CKO mouse strains (PDGFRbetaCreER:Netrin4^-/-^ and Myh11CreER:Netrin4^-/-^). It is noteworthy that the proportion of neurons responsible for intracranial innervation at the ICN terminal significantly decreased only at CKO 5 days, primarily attributable to the reduced number of neurons labeled in the periphery (**Fig 6** **D, E**). Analysis of SCG slices stained for tyrosine hydroxylase (TH) and pS6, marking neurons and protein translation activity, respectively, unveiled that, in contrast to littermate control mice, SMACreER:Netrin4^-/-^ mice exhibited a substantial decrease in the proportion of pS6-positive neurons at CKO 4 days and a significant increase in the proportion of vacuolated neurons at CKO 7 days (*P<0.05,* **Fig 6** **F-H**).

**Figure 6.**
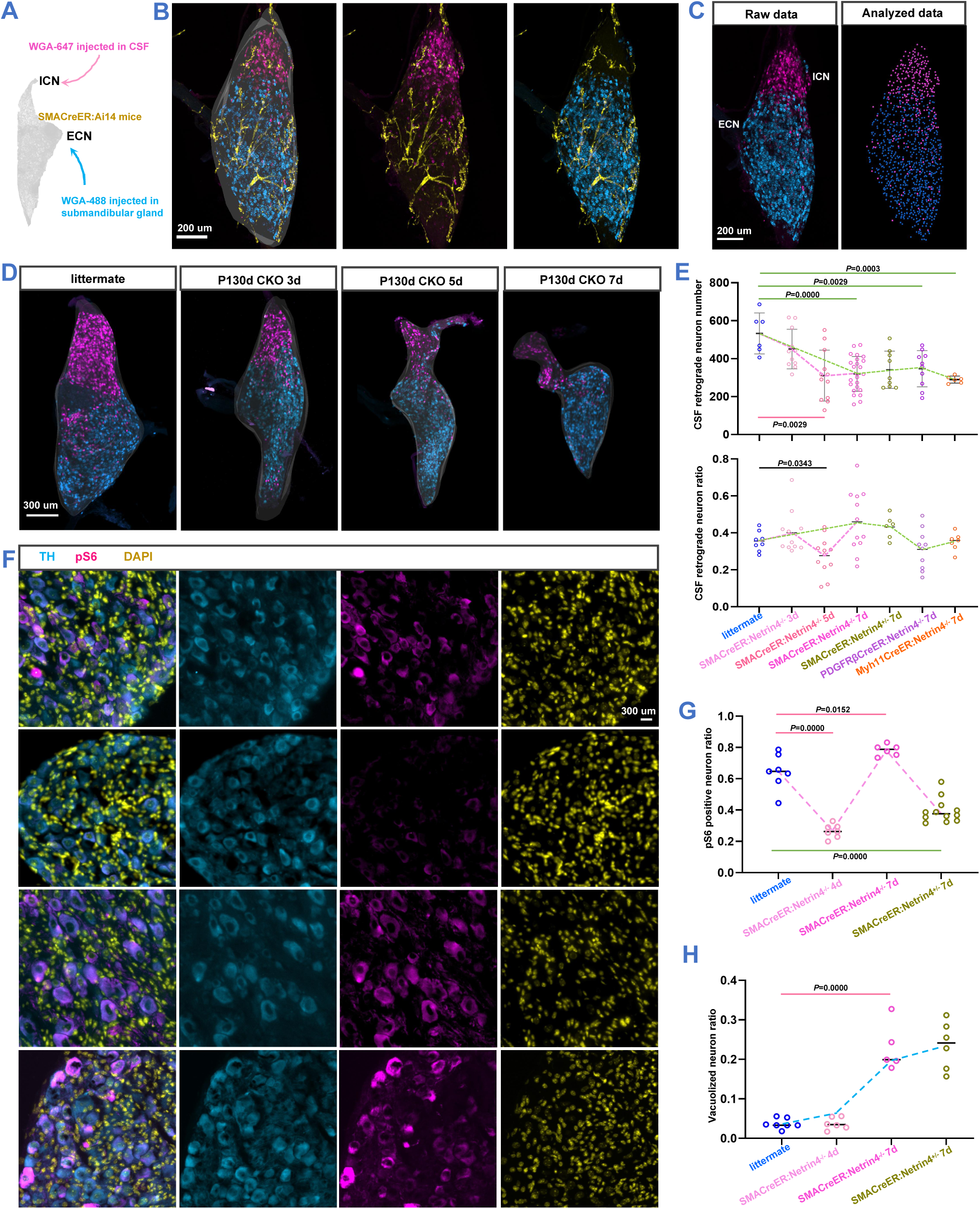
Conditional knockout of vSMCs-derived Netrin4 leads to the loss of adult mouse SCG ICN terminal neurons. **A, B, C)** Using retrograde labeling via the medullary cistern in SMACreER:Ai14 mice, WGA-Alexor647 was injected into the cerebrospinal fluid to label neurons innervating the intracranial SCG, while WGA-Alexor488 was injected into the salivary gland to label neurons innervating peripheral regions. Neuronal fluorescence signals were reconstructed as spots and counted. **D, E)** The experiment revealed that, compared to littermate control mice, the number of neurons innervating the ICN terminal in SMACreER:Netrin4^-/-^ mice at 130 days post-birth significantly decreased at CKO 5 days and 7 days. This phenomenon was also observed in heterozygotes and two other CKO mouse strains (PDGFRbetaCreER:Netrin4^-/-^ and Myh11CreER:Netrin4^-/-^). It is noteworthy that the proportion of neurons responsible for intracranial innervation at the ICN terminal significantly decreased only at CKO 5 days, mainly due to the reduced number of neurons labeled in the periphery. **F, G, H)** Analysis of SCG slices stained for TH and pS6, marking neurons and protein translation activity, respectively, revealed that, compared to littermate control mice, SMACreER:Netrin4^-/-^ mice showed a significant decrease in the proportion of pS6-positive neurons at CKO 4 days and a significant increase in the proportion of vacuolated neurons at CKO 7 days.

### 4 Netrin4 retrogradely transported to neuronal body colocalized with ribosomes

In THCreER:Ai14 mice, subsequent to the CSF injection of Netrin4-Alexor488, Netrin4-Alexor488-positive signals were observed in SCG neurons 24 hours later, with approximately 85% of Netrin4-Alexor488-positive neurons located at the ICN terminal (**Fig 7** **A-C**). Immunofluorescence staining demonstrated that Netrin4-Alexor488 signals, retrogradely transmitted from the cerebrospinal fluid, were predominantly distributed in neurons and colocalized with pS6 (**Fig 7** **D-G**). In primary cultures of adult SCG neurons treated with Netrin4 in combination with inhibitors related to endocytosis and retrograde transport, a noteworthy reduction in SCG neuron density was observed compared to the group treated with Netrin4 alone (**Fig 7** **H-J**). When WGA-Alexor555 and Netrin4-Alexor488 were simultaneously added to cultured adult SCG neurons for 15 minutes after 24 hours, WGA mainly localized to the cell membrane surface, while Netrin4 was located in the cell body (**Fig 7** **K, L**). A striking 98.7% of Netrin4-positive cells were also TH-positive cells, indicative of neurons.

**Figure 7.**
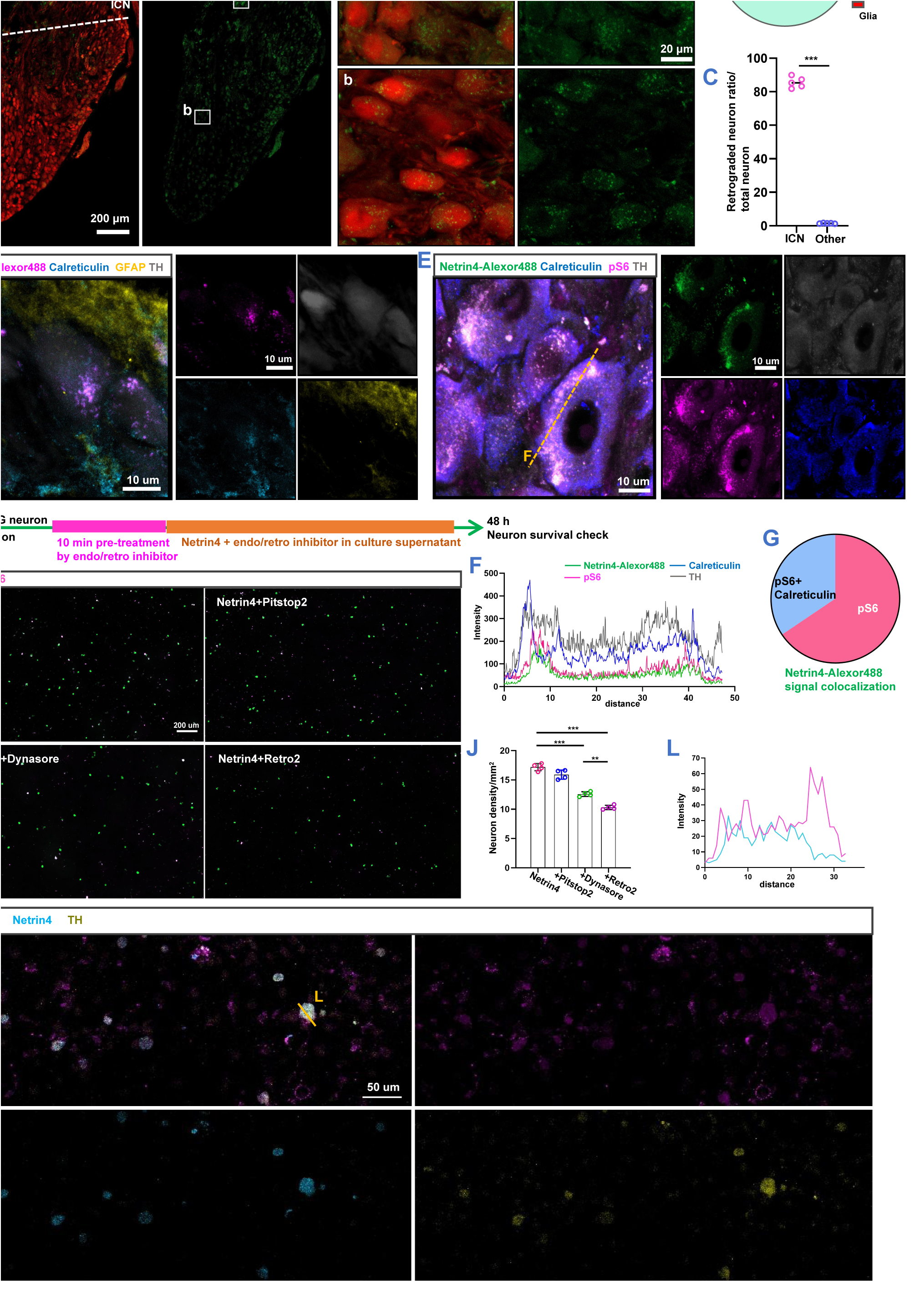
Netrin4 taken up at the axon terminals is transported retrogradely to the neuronal cell bodies and colocalizes with ribosomes. **A, B, C)** In THCreER:Ai14 mice, after cerebrospinal fluid injection of Netrin4-Alexor488, Netrin4-Alexor488-positive signals were observed in SCG neurons 24 hours later, with approximately 85% of Netrin4-Alexor488-positive neurons located at the ICN terminal. **D, E, F, G)** Immunofluorescence staining revealed that Netrin4-Alexor488 signals retrogradely transmitted from the cerebrospinal fluid were mainly distributed in neurons and colocalized with pS6. **H, I, J)** In primary cultures of adult SCG neurons treated with Netrin4 in combination with inhibitors related to endocytosis and retrograde transport, a significant reduction in SCG neuron density was observed compared to the group treated with Netrin4 alone. **K, L)** When WGA-Alexor555 and Netrin4-Alexor488 were simultaneously added to cultured adult SCG neurons for 15 minutes after 24 hours, WGA mainly localized to the cell membrane surface, while Netrin4 was located in the cell body. 98.7% of Netrin4-positive cells were also TH-positive cells, indicating neurons.

Slices examined 6 hours post-microinjection of Netrin4-Alexor488 into the SCG revealed that approximately 96% of Netrin4-Alexor488-positive cells were neurons, with approximately 85% of these neurons exhibiting strong positive pS6 staining. The immunofluorescence intensity of Netrin4-Alexor488 and pS6 in neurons displayed a positive correlation (**Fig 8** **A-D**). Following the injection of Netrin4 and Netrin4 combined with Retro2 into the SCG for 12 hours, in comparison to the contralateral side, Netrin4 injection resulted in a significant increase in ERK, Tuj1, and pS6 levels in SCG tissue, while injecting a mixture of Netrin4 and Retro2 only increased the pERK/ERK ratio (**Fig 8** **E-G**). Injection of Netrin4 alone or in combination with Retro2 into the SCG of SMACreER:Netrin4^-/-^ mice indicated that Netrin4 injection could decelerate the loss of SCG neurons, increase the proportion of pS6-positive neurons, and inhibit the level of neuronal vacuolization. However, injecting a mixture of Netrin4 and Retro2, while mitigating neuron loss, significantly increased the level of neuronal vacuolization (**Fig 8** **H-L**). Monitoring changes in cerebral blood flow for three consecutive days after local injection of Netrin4 or Netrin4 combined with Retro2 into the SCG of SMACreER:Netrin4^-/-^ mice revealed a gradual decrease in cerebral blood flow in SMACreER:Netrin4^-/-^ mice from day 3 to day 7 after tamoxifen administration. Local injection of Netrin4 maintained cerebral blood flow at pre-KO levels on day 5 and significantly increased cerebral blood flow on day 7. However, injecting a mixture of Netrin4 and Retro2 did not alter the declining trend in cerebral blood flow on day 5 but showed an upward trend on day 7 (**Fig 8** **M**). Continuous observations were conducted for 3 days using laser speckle to monitor changes in cerebral blood flow in a SMACreER:Netrin4^-/-^ mouse induced with tamoxifen for 3 days, with Netrin4 injected into the right SCG and a mixture of Netrin4 and Retro2 injected into the left SCG (**Fig 8** **N-Q**).

**Figure 8.**
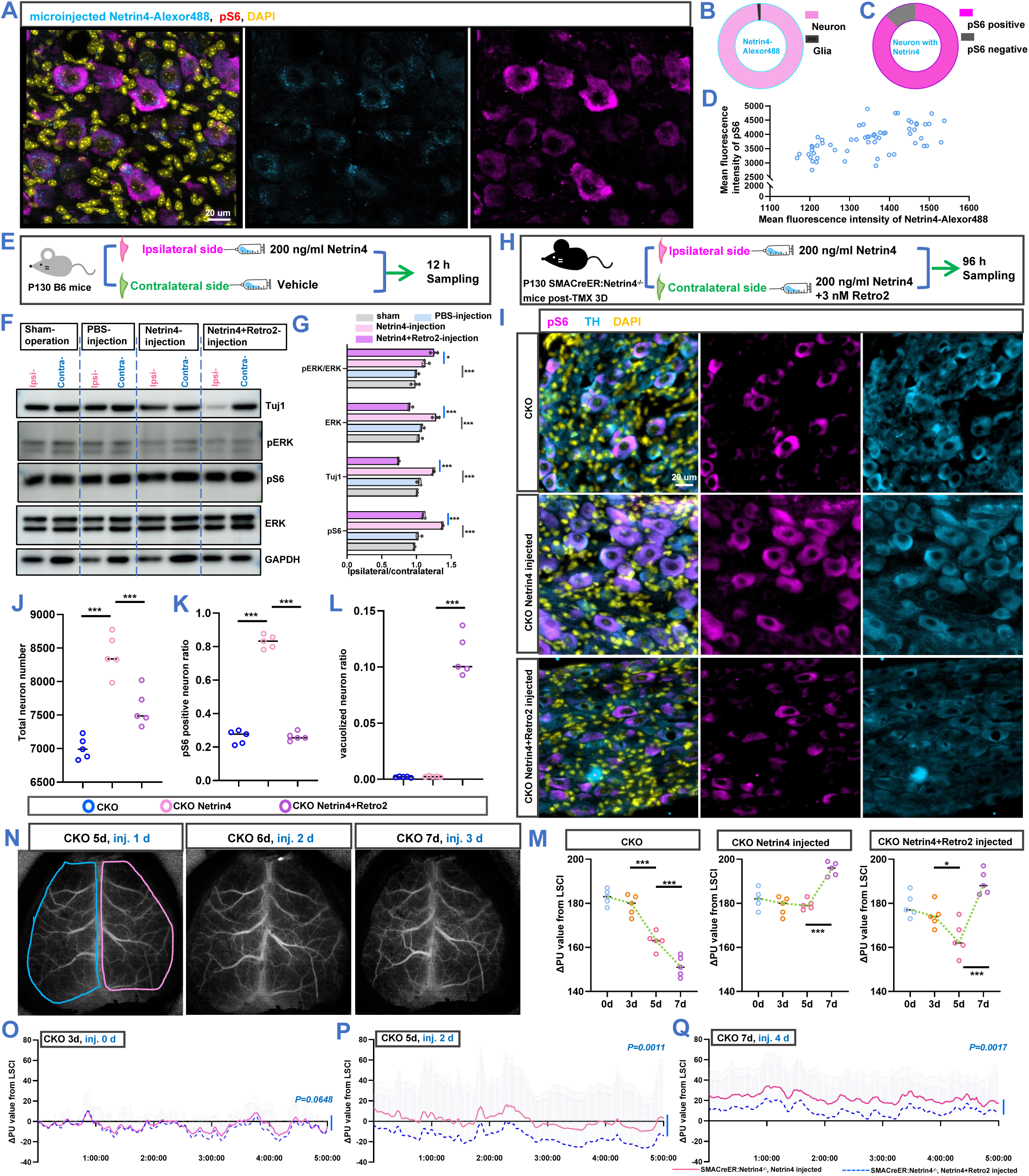
Local microinjection of Netrin4 in the SCG promotes neuronal protein synthesis and slows down the loss of SCG neurons. **A, B, C, D)** Slices observed 6 hours after microinjection of Netrin4-Alexor488 into the SCG revealed that approximately 96% of Netrin4-Alexor488-positive cells were neurons, with around 85% of these neurons showing strong positive pS6 staining. The immunofluorescence intensity of Netrin4-Alexor488 and pS6 in neurons showed a positive correlation. **E, F, G)** After injecting Netrin4 and Netrin4 combined with Retro2 into the SCG for 12 hours, compared to the contralateral side, Netrin4 injection caused a significant increase in ERK, Tuj1, and pS6 levels in SCG tissue, while injecting a mixture of Netrin4 and Retro2 only increased the pERK/ERK ratio. **I, J, K, L)** Injection of Netrin4 alone or in combination with Retro2 into the SCG of SMACreER:Netrin4^-/-^ mice showed that Netrin4 injection could slow down the loss of SCG neurons, increase the proportion of pS6-positive neurons, and inhibit the level of neuronal vacuolization. However, injecting a mixture of Netrin4 and Retro2, while alleviating neuron loss, significantly increased the level of neuronal vacuolization. **M)** Observing changes in cerebral blood flow for three consecutive days after local injection of Netrin4 or Netrin4 combined with Retro2 into the SCG of SMACreER:Netrin4^-/-^ mice revealed that cerebral blood flow gradually decreased in SMACreER:Netrin4^-/-^ mice from day 3 to day 7 after tamoxifen administration. Local injection of Netrin4 maintained cerebral blood flow at pre-KO levels on day 5 and significantly increased cerebral blood flow on day 7. However, injecting a mixture of Netrin4 and Retro2 did not alter the declining trend in cerebral blood flow on day 5 but showed an upward trend on day 7. **N, O, P, Q)** Demonstrating a SMACreER:Netrin4 CKO mouse induced with tamoxifen for 3 days, with Netrin4 injected into the right SCG and a mixture of Netrin4 and Retro2 injected into the left SCG, continuous observations were made for 3 days using laser speckle to monitor changes in cerebral blood flow.

### 5 Netrin4 sustained the levels of proteins associated with neuronal survival

Employing OPP to assess protein synthesis levels in cultured adult SCG neurons revealed that OPP-positive neurons concurrently upregulated the expression of pS6. OPP fluorescence intensity also detected protein synthesis in glial cells (**Fig 9** **A**). When using OPP to evaluate protein synthesis in mouse SCG neurons, two distinct types of OPP-positive signals were observed. One type was characterized by centrally intense staining of OPP signals in wild-type SCG neurons (**Fig 9** **B-b**). The other type exhibited a diffuse and strong OPP signal within the neurons of the ICN portion (**Fig 9** **B-a)**. Upon adding Netrin4 or a mixture of Netrin4 and Retro2 to the supernatant of adult SCG neuron cultures for 1 hour after 48 hours of incubation, compared to the neuronal basal culture medium group, Netrin4 significantly upregulated neuronal OPP levels, while the Netrin4 and Retro2 mixture only increased the fluorescence intensity of OPP in glial cells (**Fig 9** **C, D**). TH, as the key rate-limiting enzyme in neurotransmitter synthesis for sympathetic neurons in the SCG, exhibited significantly higher activity in the Netrin4-added group compared to the basal culture medium group. In contrast, the group treated with a mixture of Netrin4 and Retro2 showed a significantly reduced TH activity in neurons compared to the Netrin4-treated group (**Fig 9** **E**). Using SMACreER:Netrin4^-/-^ mice induced with tamoxifen for 4 days and 7 days to assess changes in protein synthesis levels in the SCG, it was found that, compared to littermate control mice, the SCG OPP fluorescence intensity significantly increased at CKO 7 days (**Fig 9** **F, G**). Further analysis of the density of OPP strongly positive neurons revealed a significant increase in density at CKO 4 days and 7 days in SMACreER:Netrin4^-/-^ mice. Additionally, at tamoxifen-induced 7 days, the density of OPP strongly positive neurons in heterozygous mice was also higher than that in littermate controls (**Fig 9** **H**). OPP strongly positive cells were mainly distributed at the ICN terminal (**Fig 9** **I**). In samples from the SCG of SMACreER:Netrin4^-/-^ mice induced with tamoxifen for 4 days, CSF retrograde tracing and OPP protein synthesis detection were conducted. It was observed that 95.3% of OPP-positive signals at the ICN terminal appeared in neurons positive for the retrograde tracer WGA, while 74.6% of OPP signals at the ECN segment were present in WGA-positive neurons (**Fig 9** **J**).

**Figure 9.**
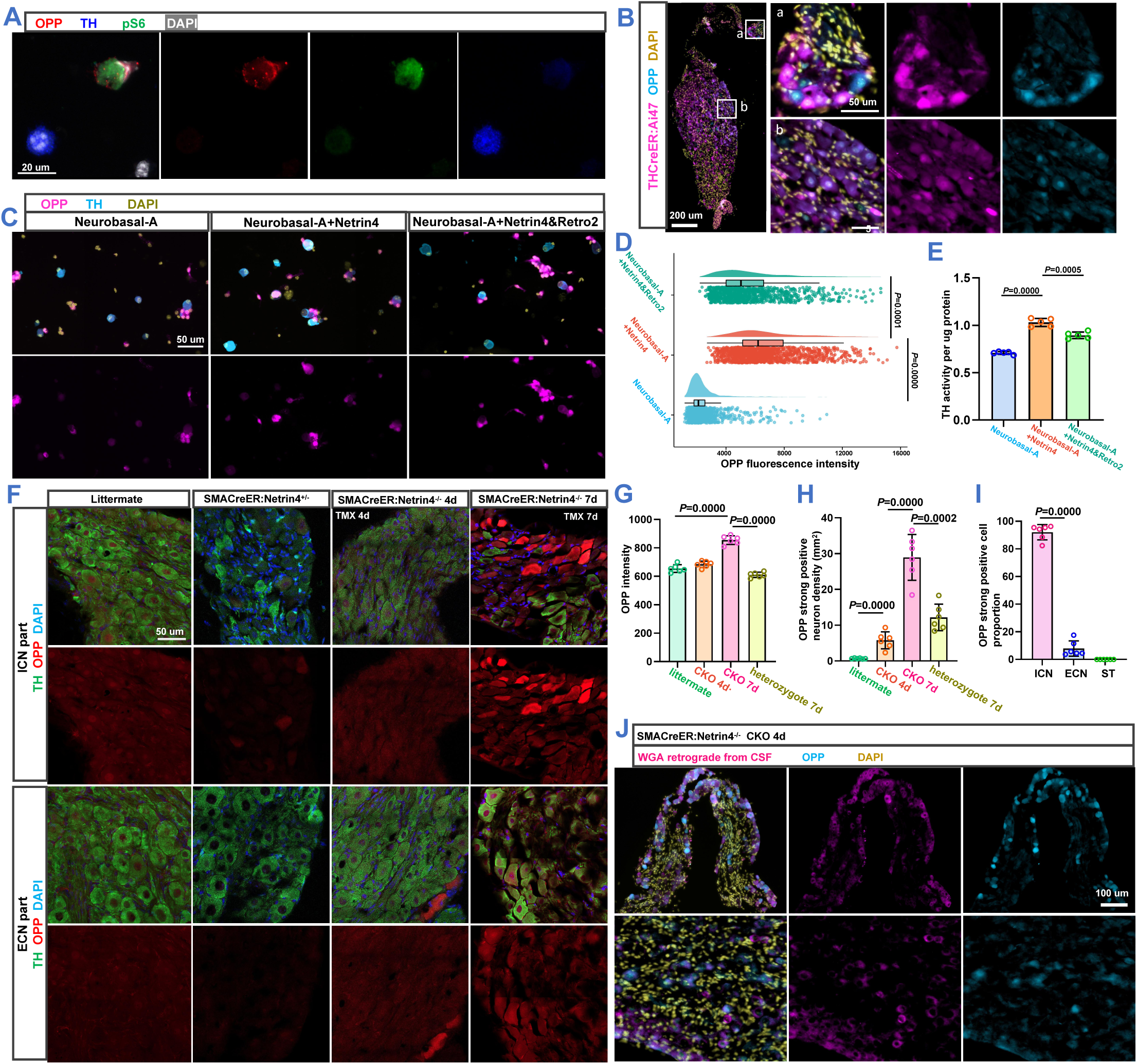
Netrin4 is internalized by neurons and maintains the levels of proteins associated with neuronal survival. **A)** Using OPP to detect protein synthesis levels in cultured adult SCG neurons, it was observed that OPP-positive neurons concurrently upregulated the expression of pS6. OPP fluorescence intensity also detected protein synthesis in glial cells. **B)** When using OPP to assess protein synthesis in mouse SCG neurons, two distinct types of OPP-positive signals were observed. One type was characterized by centrally intense staining of OPP signals in wild-type SCG neurons (b). The other type exhibited a diffuse and strong OPP signal within the neurons of ICN portion (a). **C, D, E)** When Netrin4 or a mixture of Netrin4 and Retro2 was added to the supernatant of adult SCG neuron cultures for 1 hour after 48 hours of incubation, compared to the neuronal basal culture medium group, Netrin4 significantly upregulated neuronal OPP levels, while the Netrin4 and Retro2 mixture only increased the fluorescence intensity of OPP in glial cells (D). TH, as the key rate-limiting enzyme in neurotransmitter synthesis for sympathetic neurons in the SCG, exhibits significantly higher activity in the Netrin4-added group compared to the basal culture medium group. In contrast, the group treated with a mixture of Netrin4 and Retro2 shows a significantly reduced TH activity in neurons compared to the Netrin4-treated group (E). **F, G, H, I)** Using SMACreER:Netrin4^-/-^ mice induced with tamoxifen for 4 days and 7 days to assess changes in protein synthesis levels in the SCG, it was found that, compared to littermate control mice, the SCG OPP fluorescence intensity significantly increased at CKO 7 days (G). Further analysis of the density of OPP strongly positive neurons revealed a significant increase in density at CKO 4 days and 7 days in SMACreER:Netrin4^-/-^ mice. Additionally, at tamoxifen-induced 7 days, the density of OPP strongly positive neurons in heterozygous mice was also higher than that in littermate controls (H). OPP strongly positive cells were mainly distributed at the ICN terminal (I). **J)** In samples from the SCG of SMACreER:Netrin4^-/-^ mice induced with tamoxifen for 4 days, CSF retrograde tracing and OPP protein synthesis detection were conducted. It was observed that 95.3% of OPP-positive signals at the ICN terminal appeared in neurons positive for the retrograde tracer WGA, while 74.6% of OPP signals at the ECN terminal were present in WGA-positive neurons. TH: Tyrosine hydroxylase; OPP: O-propargyl-puromycin; CSF: cerebrospinal fluid.

## Discussion

Neurons employ drastically different mechanisms to counter cell death at various stages of life[1]. The apoptotic pathways in developing neurons are highly active, aiming to finely tune the number of neurons required for the precise formation of neural networks[18]. Among neurons generated during embryonic development, over 50% disappear due to apoptosis in subsequent developmental processes[2]. In stark contrast, during adulthood, the number of neurons remains stable. Pioneering scientists, exemplified by Levi-Montalcini, discovered that SCG neurons rely on various classical neurotrophic factors such as NGF, NT3, and GDNF for their survival during the developmental process[19–22]. However, as neurons complete early development and mature, the regulatory balance between life and death undergoes a shift. Mature SCG neurons no longer rely on NGF for survival.

Numerous studies indicate that mature neurons possess a stronger ability to resist apoptosis compared to immature neurons[4]. Considering that neurons are post-mitotic cells with limited regenerative potential and may face various stresses during their lifespan, it is widely believed that mature neurons have evolved multiple, even redundant, mechanisms to rigorously prevent apoptosis, ensuring their long-term survival.

However, mature neurons throughout their lifespan constantly face microenvironmental pressures. In addition to programmed cell apoptosis, potential ways to induce neuronal death include necroptosis, ferroptosis, and pyroptosis, among more than a dozen other mechanisms[23–26]. While mature neurons can survive independently of well-known neurotrophic factors such as NGF, the question arises: do they still rely on other nutritional factors from their targets for survival? Does this nutritional factor selectively inhibit another death pathway independent of apoptosis? These scientific queries are crucial for understanding the survival mechanisms of mature neurons, yet there is currently no reported research on these aspects. Moreover, David D. Ginty of Johns Hopkins University, an eminent scientist in the field, believes that there may still be undiscovered nutritional factors even during the developmental process, warranting further investigation.

This study found that depriving target cells, aSMCs, of their ability to secrete Netrin-4 during adulthood (SMACreER:Ntn4^f/f^) led to a 45% decrease in the number of mature neurons in the SCG. Consequently, the volume of the ganglion decreased by 50%, and p-MLKL-positive neurons appeared in the SCG ganglion. MLKL (Mixed lineage kinase domain-like Pseudokinase) is a marker protein for programmed cell necrosis, necroptosis. This implies that the Netrin-4 protein secreted by aSMCs is a necessary condition for inhibiting programmed cell necrosis in mature SCG neurons.

Further studies in several aspects have confirmed our speculations. Firstly, by tracking SCG neurons responsible for innervating the intracranial region in CKO mice, we observed that neurons responsible for innervating the intracranial region were lost early (5 days) after CKO induction, followed by the loss of neurons responsible for innervating the periphery at 7 days, leading to a significant reduction in SCG volume. These results reveal the temporal progression of SCG neuron loss in SMACreER:Ntn4^f/f^ mice. Secondly, we found that a higher concentration of p-MLKL-positive neurons was clustered at the ICN end, where approximately 85% of neurons are responsible for innervating intracranial structures, and this area is the first to experience neuron loss. Thirdly, local supplementation of Netrin-4 in the SCG significantly alleviated the neuron loss induced by CKO, indicating that Netrin-4 is a critical factor in the loss of SCG neurons in SMACreER:Ntn4^f/f^ mice. Additionally, we characterized the volume curve of the SCG at different stages of a mouse’s life. We found that the SCG volume in mice continues to increase until 130 days after birth, reaching its maximum level, and then begins to decline at 230 days. After inducing CKO in SMACreER:Ntn4^f/f^ mice for 7 days, there was a significant reduction in SCG volume in mice aged 130-270 days, accompanied by neuron loss. These observations suggest that Netrin-4 derived from target cells may be one of the key factors in maintaining the survival of adult SCG neurons.

We observed an interesting phenomenon: whether adding Netrin-4 to the culture supernatant of SCG cells or injecting it directly into the SCG region, Netrin-4 was more often detected in the neuronal cell bodies rather than on the membrane. When we injected Netrin-4 into the CSF, it also retrogradely transported from the axonal end back to the neuronal cell bodies, co-localizing with ribosomes. This suggests that neurons are prone to internalize Netrin-4. Furthermore, when we added inhibitors of endocytosis and retrograde transport while culturing SCG neurons with Netrin-4, the neuroprotective effect of Netrin-4 was significantly reduced. This indicates that Netrin-4 needs to be internalized from the axonal end by neurons and transported to the cell body to exert its protective effect on neurons.

When we observed that local replenishment of Netrin-4 in the SCG could rescue the loss of SCG neurons in SMACreER:Ntn4^f/f^ mice, we wondered whether retrograde transport was necessary for the neuroprotective effect of Netrin-4 at this point. We injected Netrin-4 into the SCG of wild-type mice and found a significant upregulation in the protein translation level in the SCG. However, co-administration of Retro2 markedly inhibited this effect, further confirming that retrograde transport is crucial for the neuroprotective action of Netrin-4 internalized into neurons.

A series of in vivo and in vitro experiments consistently indicate that pS6 is consistently associated with the survival of SCG neurons during Netrin-4-induced processes. Moreover, the co-localization of Netrin-4 with ribosomes after internalization into neurons further raises our suspicion that it may be directly related to protein synthesis.

OPP was utilized in both in vivo and in vitro experiments to validate whether Netrin-4 can influence the protein synthesis levels of neurons. In our findings, the addition of Netrin-4 to the culture supernatant of mature SCG neurons in vitro significantly upregulated the levels of OPP in neurons, while the levels in glial cells remained unchanged. This suggests that Netrin-4 can enhance the protein synthesis levels in SCG neurons. However, in SMACreER:Ntn4^f/f^ mice during the CKO process, we observed strong OPP-positive signals in neurons at the ICN end. Through retrograde tracing, we found that these cells were predominantly responsible for innervating the intracranial region. This indicates that the observed protein synthesis signal during in vivo CKO is mediated by neurons lacking Netrin-4, and the proteins synthesized under Netrin-4 deprivation in vivo may differ significantly from those increased by the addition of Netrin-4 in in vitro experiments. This distinction plays a crucial role in determining the life or death of SCG neurons.

The elucidation of downstream proteins crucial for maintaining the survival of mature SCG neurons by the target organ-secreted Netrin-4 will be a focal point in our upcoming experiments. These proteins may serve as brakes for necroptosis in mature SCG neurons and represent key points in the reshaping of the SCG induced directly by the activity and functional status of the target organ secretion. The clarification of the mechanism by which Netrin-4 maintains the survival of mature neurons could potentially lead to the identification of the first neuroprotective factor directly combating neuronal necroptosis.

## Acknowledgements

We thank Dongdong Zhang, Jinze Li, Tingbo Li for their constructive discussions. We thank Danping Lu for her responsive and timely purchasing support. We thank the animal facility for its technical assistance with rodent housing, Biomedical Research center platform for technique support. This work is supported by the National Natural Science Foundation of China (31800864 and 31970969 to J.-M.J, 82101475 to Z.Z.), Westlake startup funds, Westlake Education Foundation and MRIC funds (103536022011) to J.-M.J.

## Author contributions

J.-M.J. conceived the research and designed experiments. Z.Z. performed all the experiments and data quantification. L.L.Z conducted the bulk RNA sequence data analysis. Y.J.W supplied the mouse management. Y.J.W and X.X.Z conducted the neuron number analysis. Dongdong Zhang gave precise protocol for immunofluorescence. Xuzhao Li designed the transgenetic mice. Bingrui Zhao conducted mouse genotyping.

---

**Figure S1.**
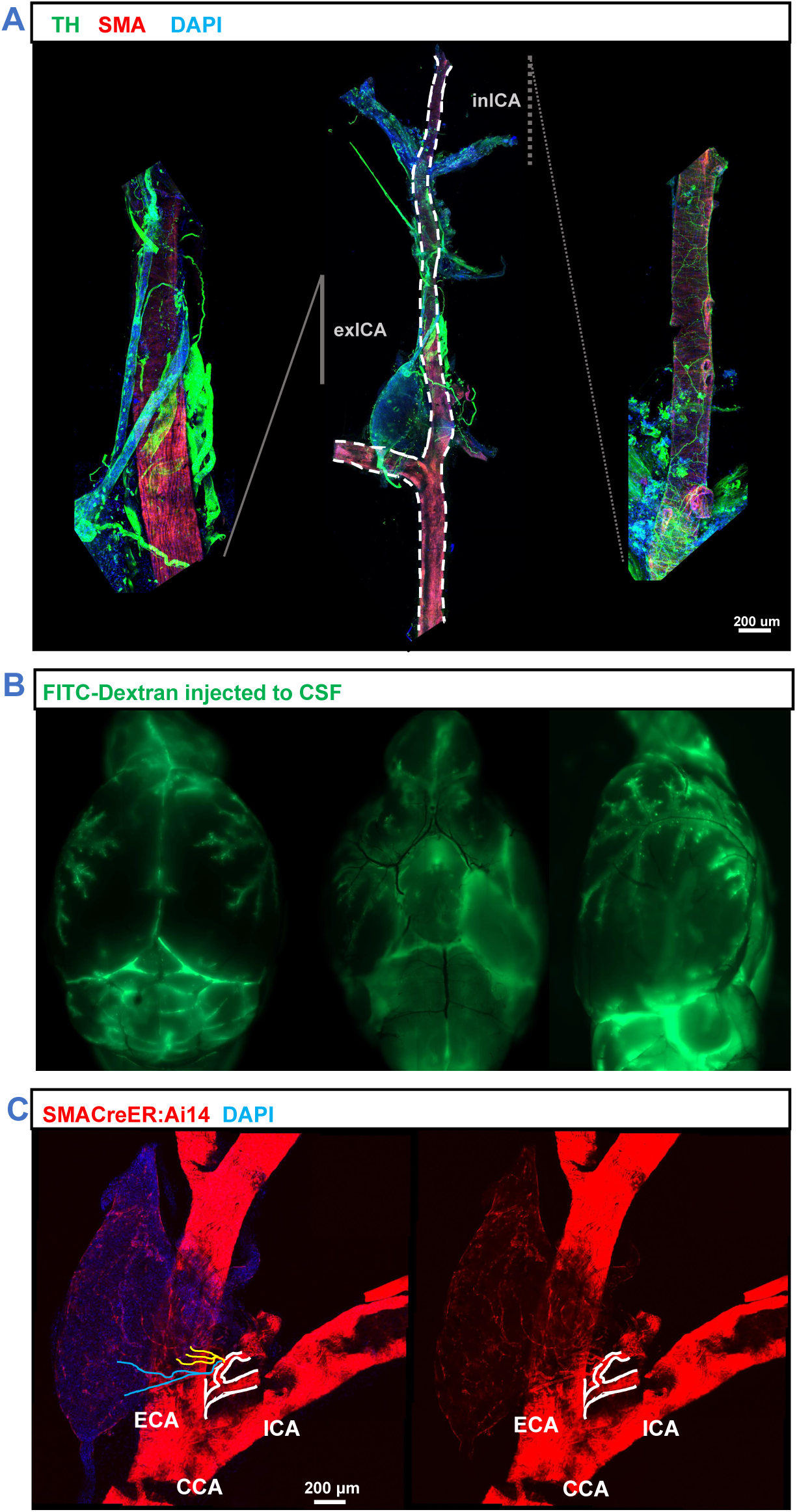
Morphological Study of the Vascular Supply and Postganglionic Fiber Distribution in the SCG. **A)** The main branches of the CCA and ICA were isolated. Immunofluorescent staining was conducted using tyrosine hydroxylase (TH) to label postganglionic fibers of the sympathetic nervous system, while smooth muscle cells of blood vessels were marked with smooth muscle actin (SMA). Postganglionic nerve fibers of the SCG formed bundles ascending along the ICA, and upon entering the cranial cavity through the carotid foramen, they exhibited a plexiform arrangement surrounding the ICA. **B)** Dextran-FITC was injected into the cerebellomedullary cistern of mice, and after 20 minutes, the brain was extracted. The FITC fluorescent signal was predominantly observed in the perivascular space of the meningeal arteries. **C)** Tissue samples were obtained from the bifurcation of the CCA and the SCG in SMACreER:Ai14 mice. The primary blood supply arteries to the SCG originated from the branches of the ECA, specifically the lingual artery. Immediately upon branching, the lingual artery split into two branches, both entering the SCG at the ECN end. The upper branch further divided into three main branches, supplying the ICN end and ECN end of the SCG, while the lower branch was responsible for supplying the ST segment.

**Figure S2.**
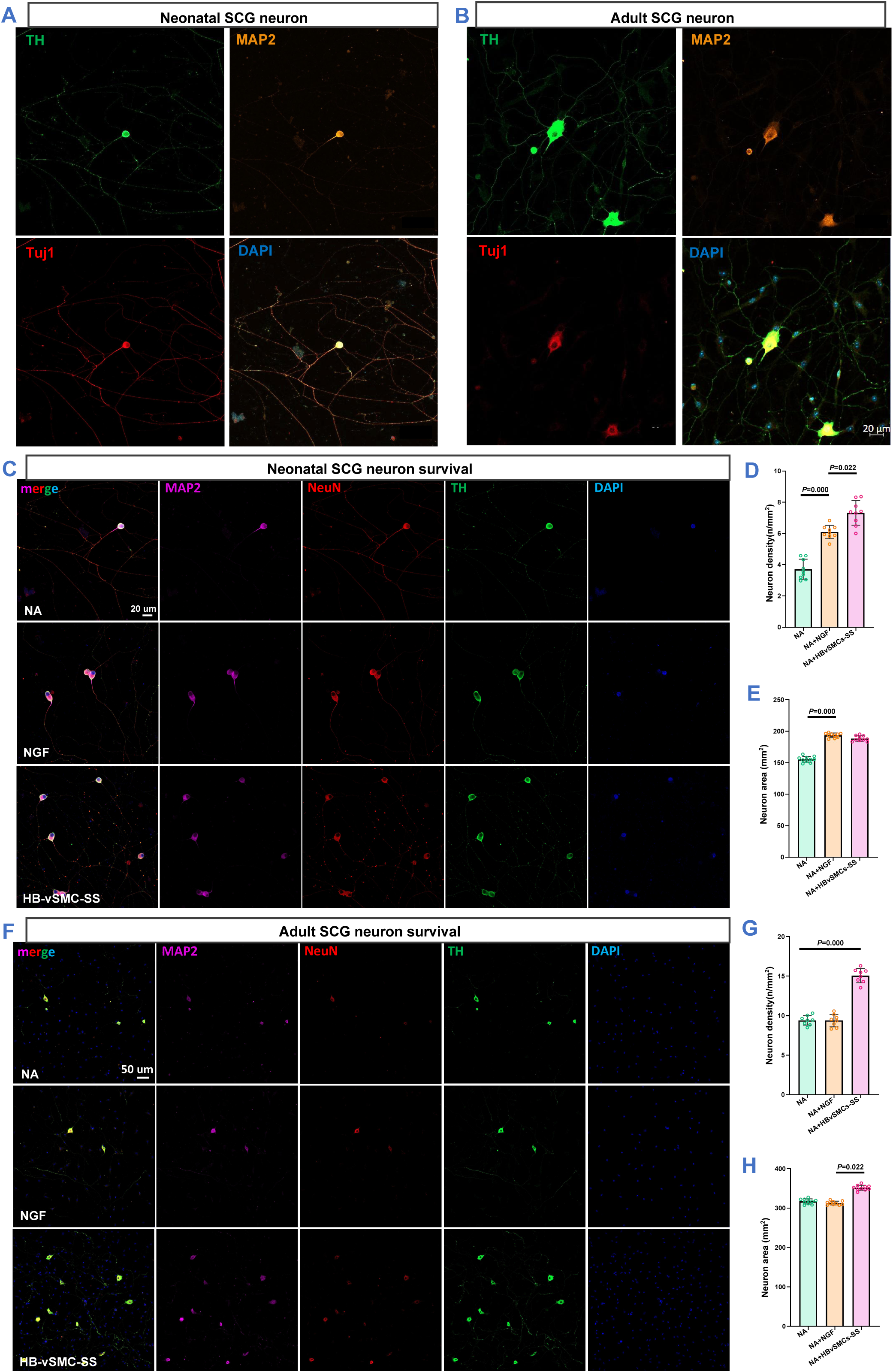
**The secreted supernatant** from human brain-derived smooth muscle cells promotes the survival of neonatal and adult SCG neurons. **A, C, D, E)** Primary culture of mouse SCG neurons from postnatal day 0 to 1. NGF significantly enhances the survival of neonatal SCG neurons, while the secreted supernatant from HBvSMCs markedly promotes the survival of neonatal SCG neurons, with its effectiveness surpassing that of NGF. NGF and the secreted supernatant from HBvSMCs significantly increase the size of cultured neurons compared to the basal culture medium. **B, F, G, H)** Primary culture of mouse SCG neurons from postnatal day 60. NGF does not alter the survival levels in the in vitro culture of adult SCG neurons, whereas the conditioned medium from HBvSMCs significantly enhances the size and survival of adult SCG neurons.

**Figure S3.**
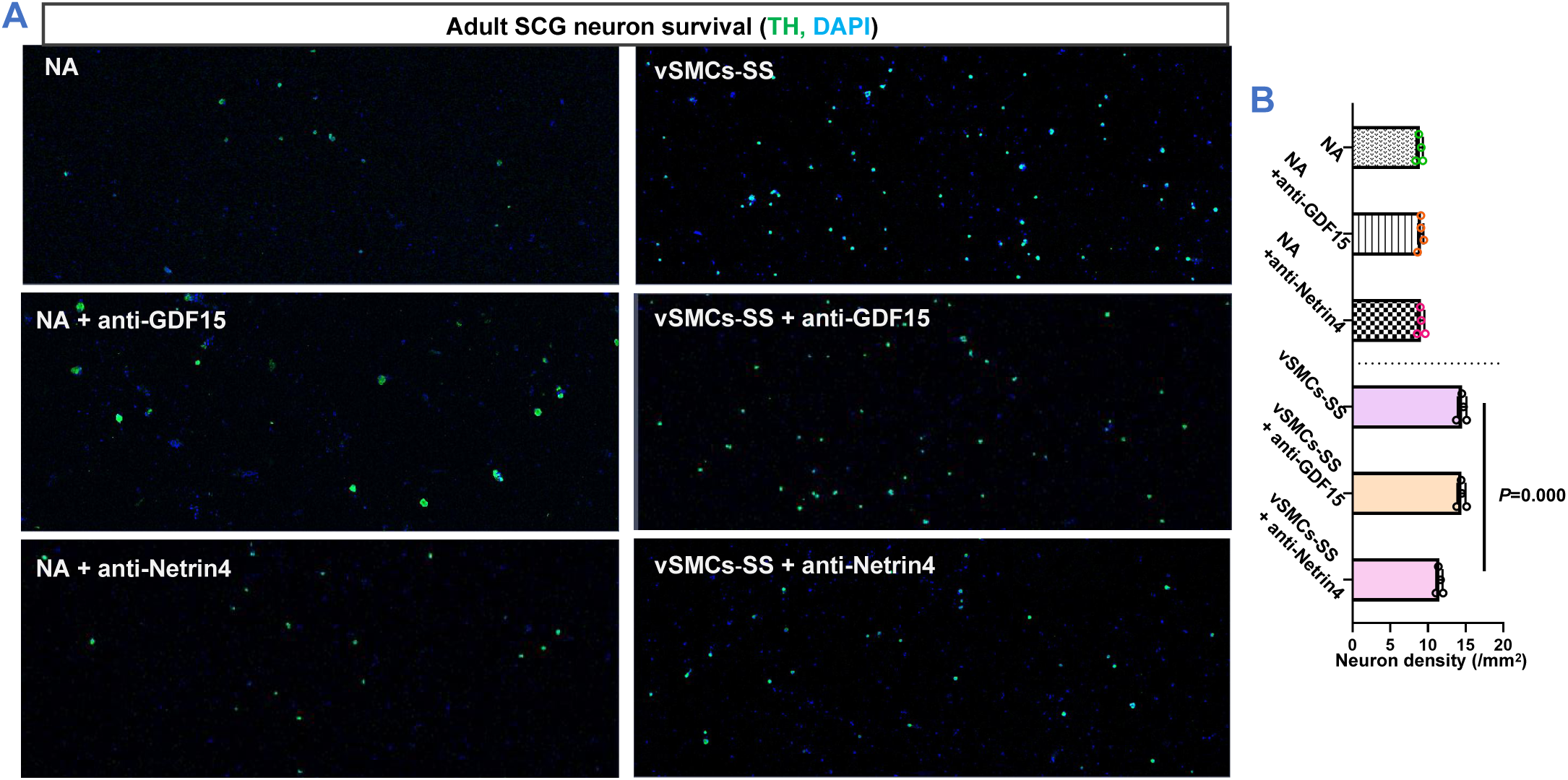
Adult neuron protective candidate selection from vSMCs secretome. After separately neutralizing GDF-15 and Netrin4 in the secreted supernatant of cerebral vSMCs with antibodies, it was observed that the survival density of adult SCG neurons cultured in the supernatant with added Netrin4 antibody was significantly lower than that in the vSMCs supernatant culture group, but higher than that in the Neurobasal-A culture group. However, neutralizing GDF-15 in the vSMCs secreted supernatant with antibodies did not impact the neuronal survival density.

**Figure S4.**
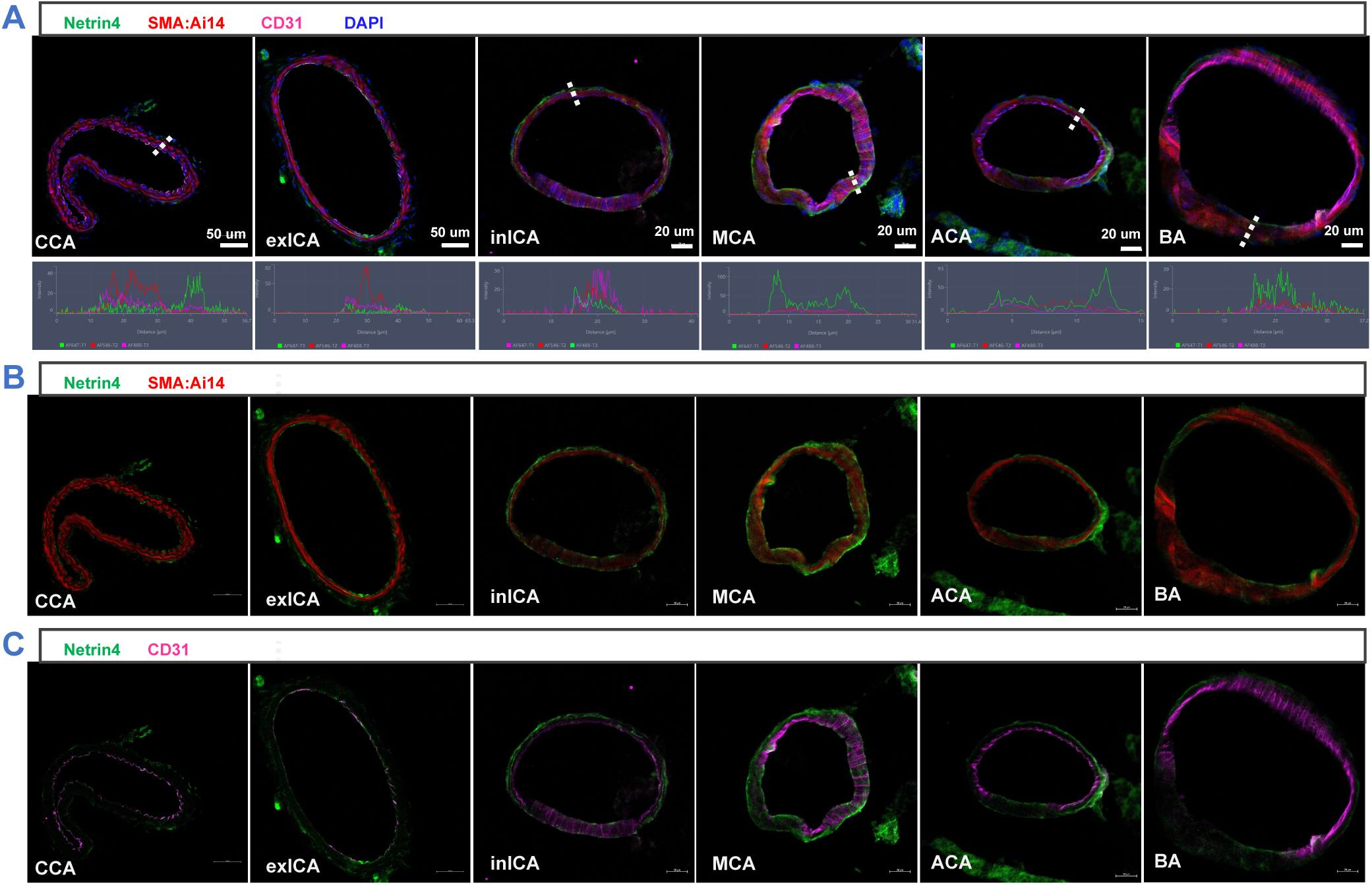
Netrin4 is distributed in intracranial arterial SMCs and the basement membrane, promoting protein synthesis in adult sympathetic neurons. **A, B, C)** Cross-sectional slices of intracranial and extracranial arteries were subjected to immunofluorescence staining, revealing that Netrin4 is predominantly distributed in intracranial arteries (A). Its primary expression occurs in vSMCs and the basement membrane (B), with no detectable Netrin4 signal in endothelial cells marked by CD31 (C).

**Figure S5.**
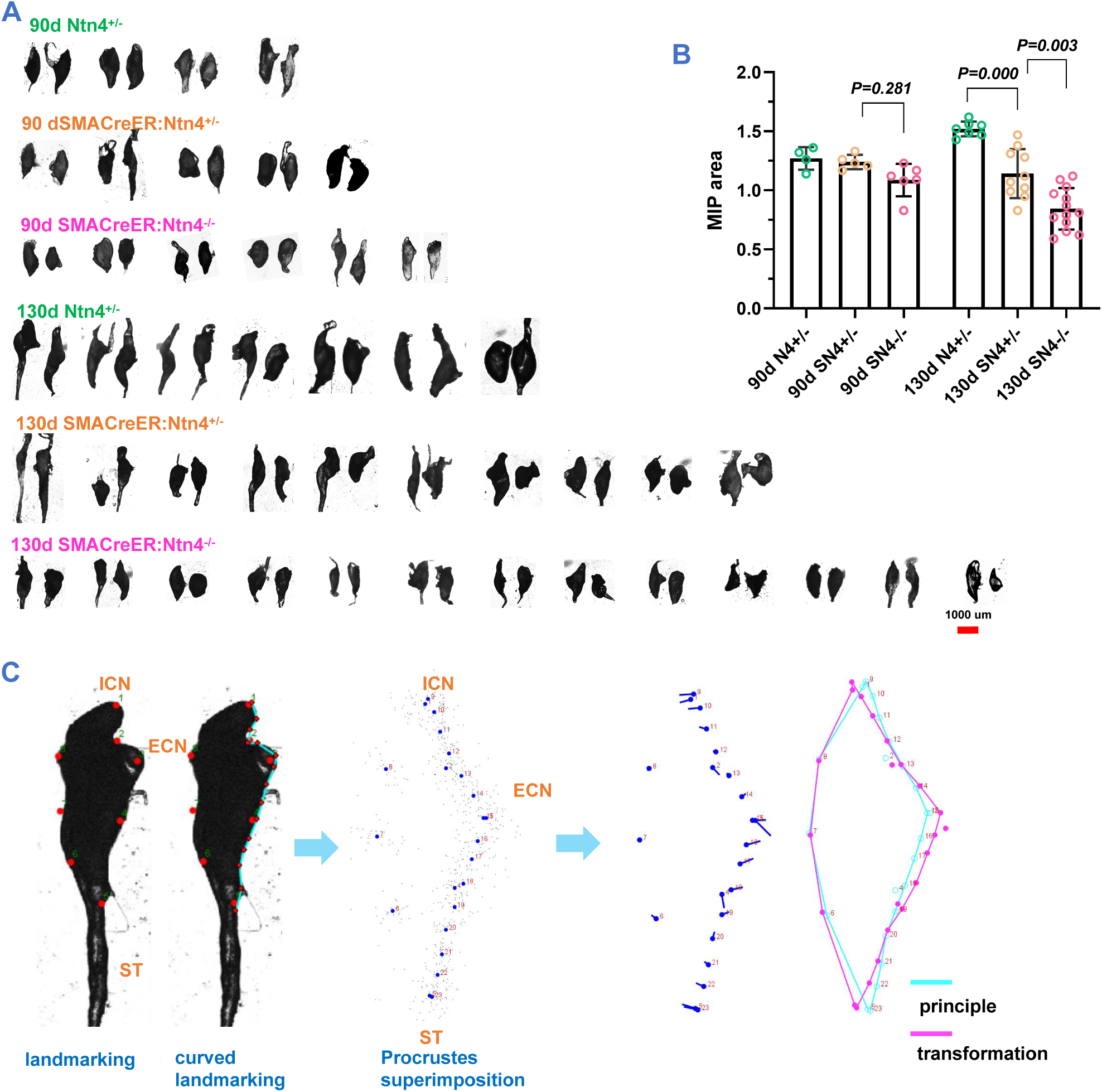
Conditional knockout of vSMCs-derived Netrin4 causes a reduction in adult mouse SCG size. **A, B)** Compared to littermate control mice, conditional knockout of arterial SMC Netrin4 in SMACreER:Netrin4 mice at 130 days post-birth resulted in a significant decrease in the size of both homozygous and heterozygous SCGs after 7 days. In contrast, conditional knockout mice at 90 days post-birth showed a reduced SCG size in homozygotes, while heterozygotes exhibited no significant changes. **C)** The standardization of SCG morphology using the Landmark method serves as the foundational approach in this study to analyze localized changes in SCG volume.

