## Supplementary material for "Netrin4 is a new target specific factor, ensuring adult sympathetic neuron survival via promoting protein synthesis": Fig S: supple Netrin4 is a new target specific factor, ensuring adult sympathetic neuron survival via promoting protein synthesis.pdf

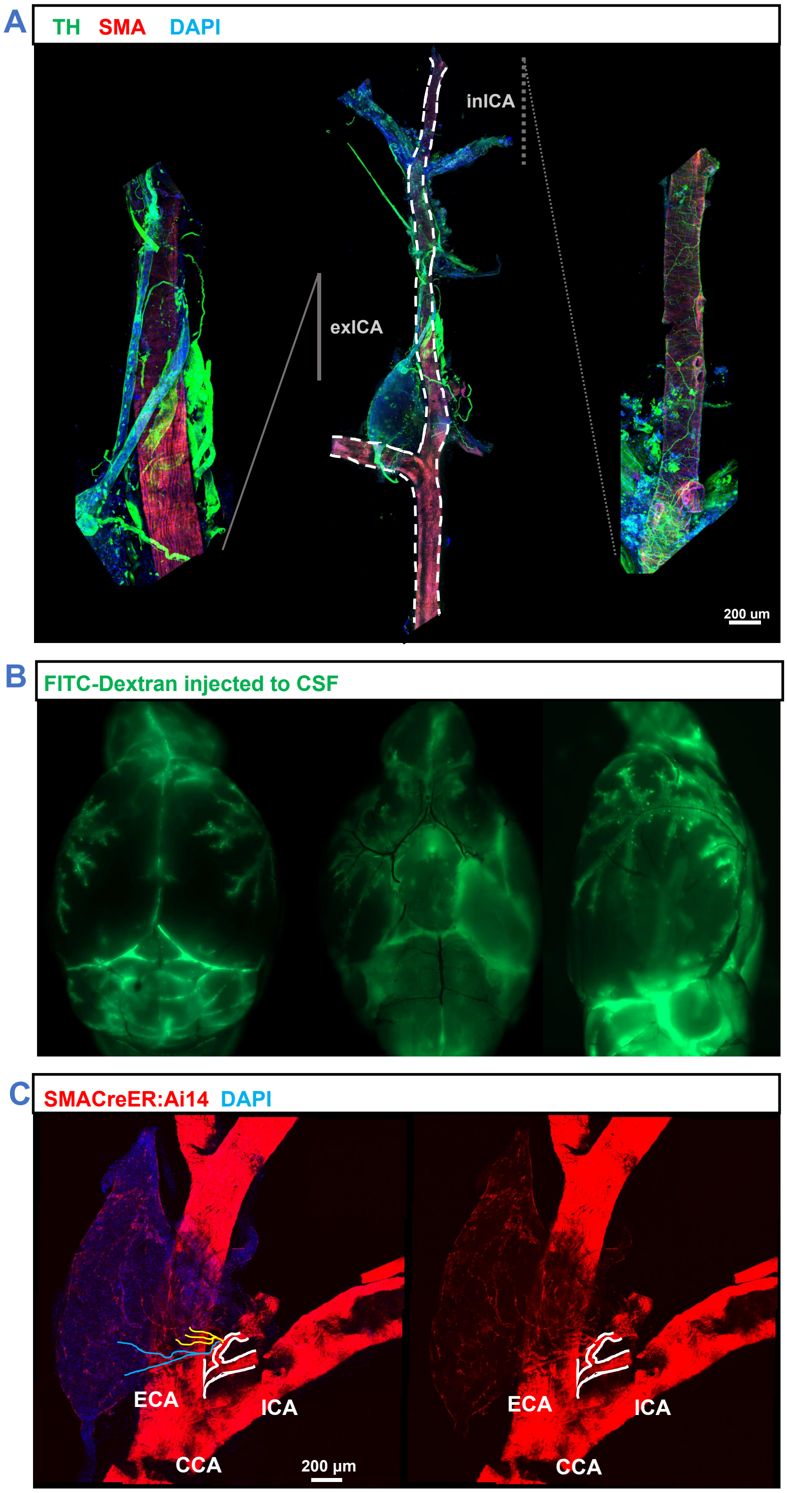

A
TH SMA DAPI
inICA
exICA
200 um
B
FITC-Dextran injected to CSF
C
SMACreER:Ai14 DAPI
ECA
ICA
CCA
200 μm
ECA
ICA
CCA

**Figure S1 Morphological Study of the Vascular Supply and Postganglionic Fiber Distribution in the SCG. A)** The main branches of the CCA and ICA were isolated. Immunofluorescent staining was conducted using tyrosine hydroxylase (TH) to label postganglionic fibers of the sympathetic nervous system, while smooth muscle cells of blood vessels were marked with smooth muscle actin (SMA). Postganglionic nerve fibers of the SCG formed bundles ascending along the ICA, and upon entering the cranial cavity through the carotid foramen, they exhibited a plexiform arrangement surrounding the ICA. **B)** Dextran-FITC was injected into the cerebellomedullary cistern of mice, and after 20 minutes, the brain was extracted. The FITC fluorescent signal was predominantly observed in the perivascular space of the meningeal arteries. **C)** Tissue samples were obtained from the bifurcation of the CCA and the SCG in SMACreER:Ai14 mice. The primary blood supply arteries to the SCG originated from the branches of the ECA, specifically the lingual artery. Immediately upon branching, the lingual artery split into two branches, both entering the SCG at the ECN end. The upper branch further divided into three main branches, supplying the ICN end and ECN end of the SCG, while the lower branch was responsible for supplying the ST segment.

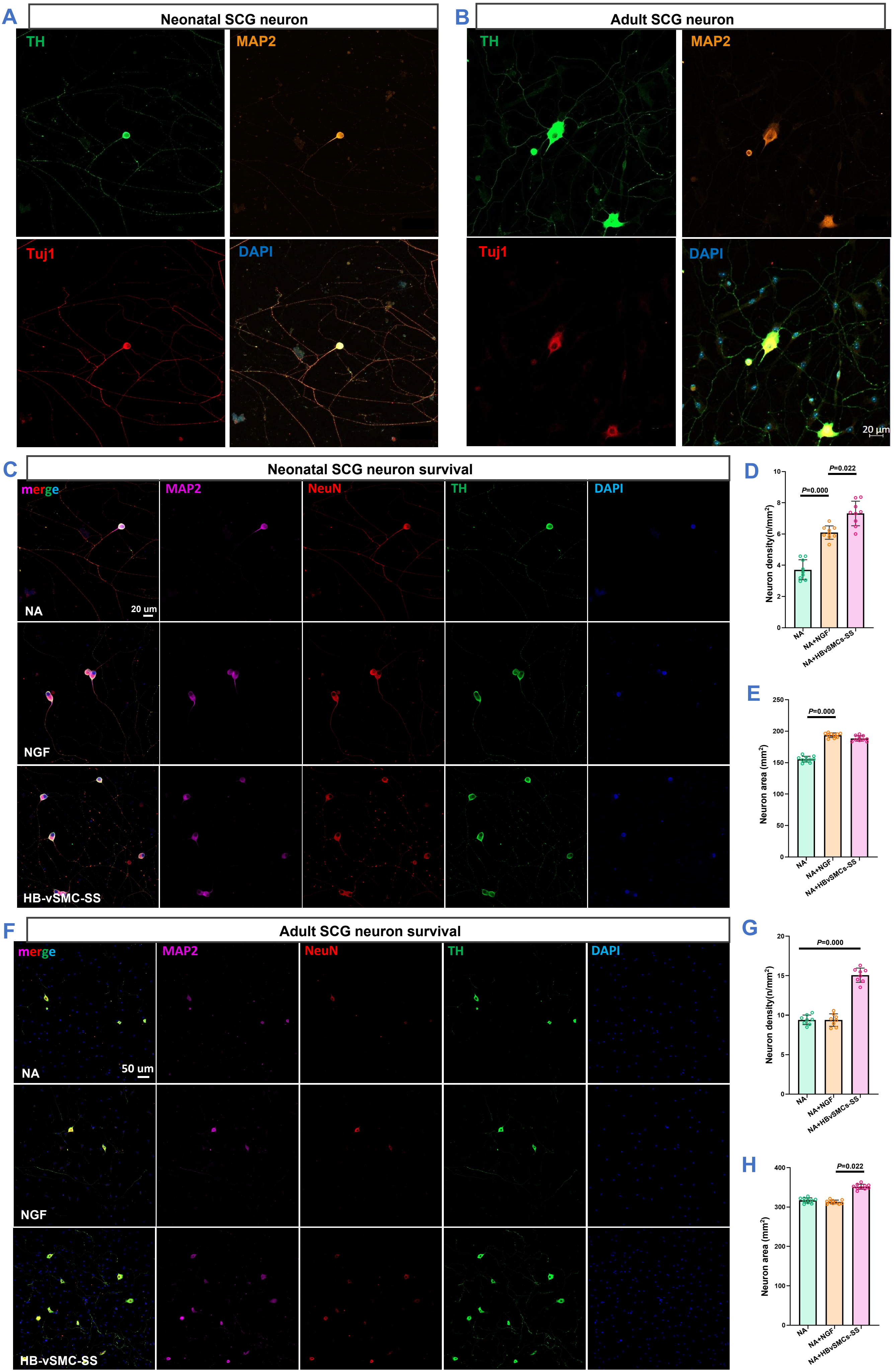

A
Neonatal SCG neuron
TH
MAP2
Tuj1
DAPI
B
Adult SCG neuron
TH
MAP2
Tuj1
DAPI
20 μm
C
Neonatal SCG neuron survival
merge
MAP2
NeuN
TH
DAPI
NA
20 um
NGF
HB-vSMC-SS
D
P=0.000
P=0.022
Neuron density(n/mm²)
NA
NA+NGF
NA+HBvSMCs-SS
E
P=0.000
Neuron area (mm²)
NA
NA+NGF
NA+HBvSMCs-SS
F
Adult SCG neuron survival
merge
MAP2
NeuN
TH
DAPI
NA
50 um
NGF
HB-vSMC-SS
G
P=0.000
Neuron density(n/mm²)
NA
NA+NGF
NA+HBvSMCs-SS
H
P=0.022
Neuron area (mm²)
NA
NA+NGF
NA+HBvSMCs-SS

**Figure S2 The secreted supernatant** from human brain-derived smooth muscle cells promotes the survival of neonatal and adult SCG neurons. **A, C, D, E)** Primary culture of mouse SCG neurons from postnatal day 0 to 1. NGF significantly enhances the survival of neonatal SCG neurons, while the secreted supernatant from HBvSMCs markedly promotes the survival of neonatal SCG neurons, with its effectiveness surpassing that of NGF. NGF and the secreted supernatant from HBvSMCs significantly increase the size of cultured neurons compared to the basal culture medium. **B, F, G, H)** Primary culture of mouse SCG neurons from postnatal day 60. NGF does not alter the survival levels in the in vitro culture of adult SCG neurons, whereas the conditioned medium from HBvSMCs significantly enhances the size and survival of adult SCG neurons.

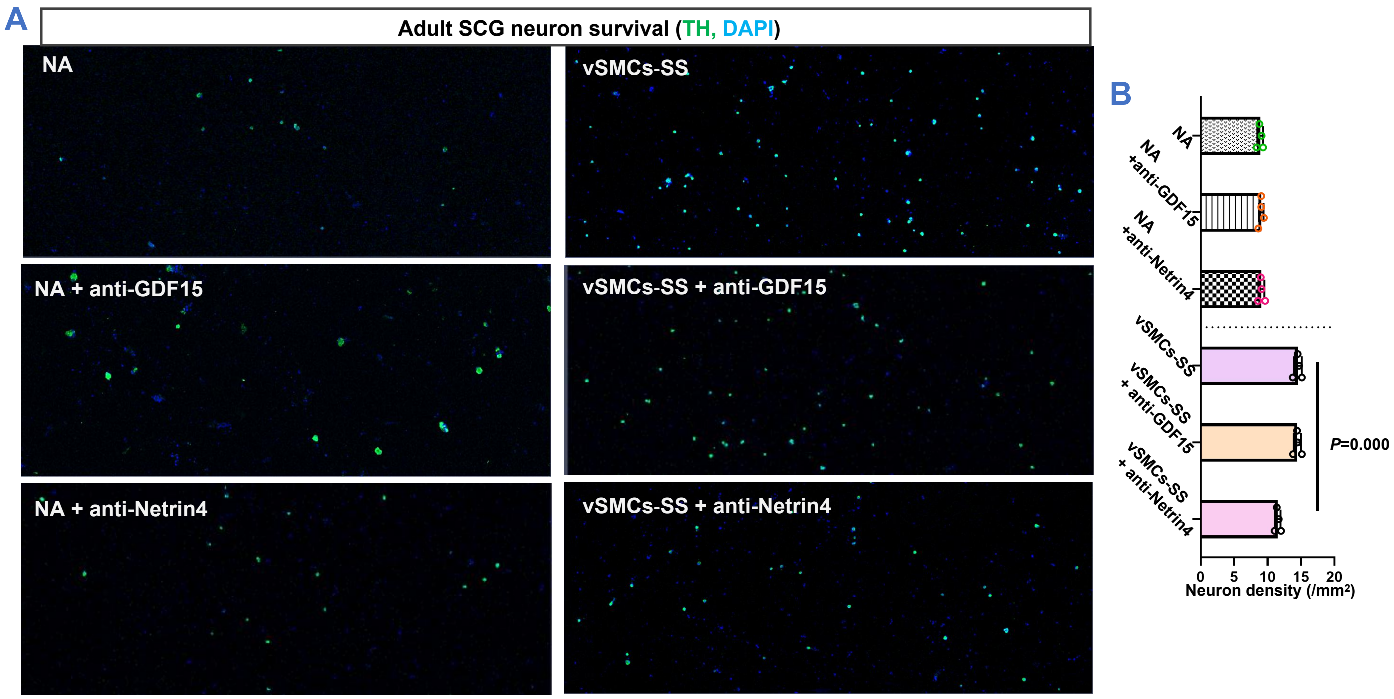
A
Adult SCG neuron survival (TH, DAPI)
NA
vSMCs-SS
NA + anti-GDF15
vSMCs-SS + anti-GDF15
NA + anti-Netrin4
vSMCs-SS + anti-Netrin4
B
NA
NA +anti-GDF15
NA +anti-Netrin4
vSMCs-SS
vSMCs-SS + anti-GDF15
vSMCs-SS + anti-Netrin4
P=0.000
0 5 10 15 20
Neuron density (/mm²)

**Figure S3 Adult neuron protective candidate selection from vSMCs secretome.** After separately neutralizing GDF-15 and Netrin4 in the secreted supernatant of cerebral vSMCs with antibodies, it was observed that the survival density of adult SCG neurons cultured in the supernatant with added Netrin4 antibody was significantly lower than that in the vSMCs supernatant culture group, but higher than that in the Neurobasal-A culture group. However, neutralizing GDF-15 in the vSMCs secreted supernatant with antibodies did not impact the neuronal survival density.

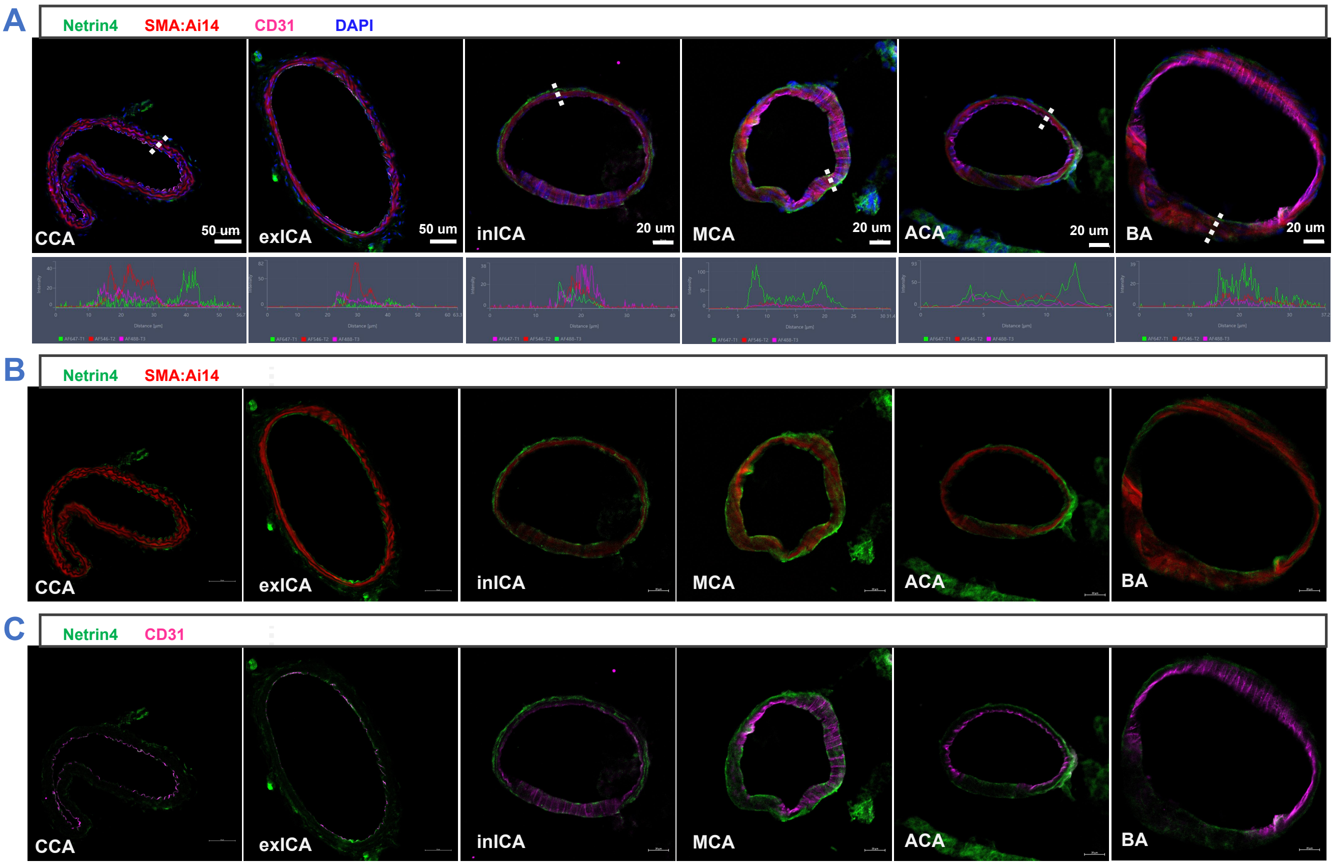
A
Netrin4
SMA:Ai14
CD31
DAPI
CCA
50 um
exICA
50 um
inICA
20 um
MCA
20 um
ACA
20 um
BA
20 um
B
Netrin4
SMA:Ai14
CCA
exICA
inICA
MCA
ACA
BA
C
Netrin4
CD31
CCA
exICA
inICA
MCA
ACA
BA

**Figure S4 Netrin4 is distributed in intracranial arterial SMCs and the basement membrane, promoting protein synthesis in adult sympathetic neurons. A, B, C)** Cross-sectional slices of intracranial and extracranial arteries were subjected to immunofluorescence staining, revealing that Netrin4 is predominantly distributed in intracranial arteries (A). Its primary expression occurs in vSMCs and the basement membrane (B), with no detectable Netrin4 signal in endothelial cells marked by CD31 (C).

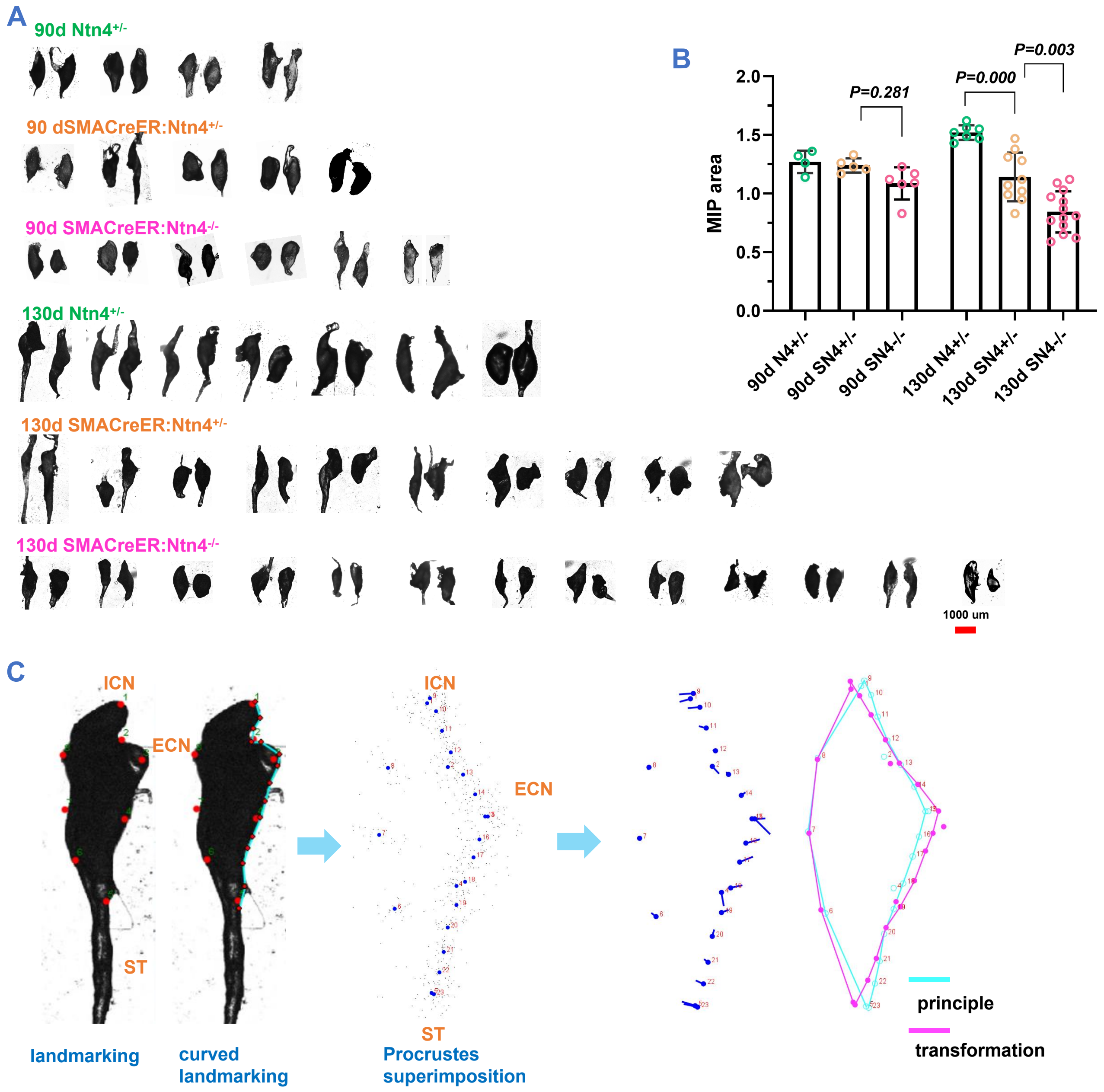

A
90d Ntn4+/-
90 dSMACreER:Ntn4+/-
90d SMACreER:Ntn4-/-
130d Ntn4+/-
130d SMACreER:Ntn4+/-
130d SMACreER:Ntn4-/-
1000 um
B
MIP area
2.0
1.5
1.0
0.5
0.0
P=0.281
P=0.000
P=0.003
90d N4+/-
90d SN4+/-
90d SN4-/-
130d N4+/-
130d SN4+/-
130d SN4-/-
C
ICN
ECN
ST
landmarking
curved landmarking
Procrustes superimposition
principle
transformation

**Figure S5 Conditional knockout of vSMCs-derived Netrin4 causes a reduction in adult mouse SCG size**. **A, B)** Compared to littermate control mice, conditional knockout of arterial SMC Netrin4 in SMACreER:Netrin4 mice at 130 days post-birth resulted in a significant decrease in the size of both homozygous and heterozygous SCGs after 7 days. In contrast, conditional knockout mice at 90 days post-birth showed a reduced SCG size in homozygotes, while heterozygotes exhibited no significant changes. **C)** The standardization of SCG morphology using the Landmark method serves as the foundational approach in this study to analyze localized changes in SCG volume.
